# A genome-wide overexpression screen reveals *Mycobacterium smegmatis* growth inhibitors encoded by mycobacteriophage Hammy

**DOI:** 10.1101/2023.06.21.545952

**Authors:** Isabel Amaya, Kaylia Edwards, Bethany M. Wise, Ankita Bhattacharyya, Clint H. D. Pablo, Ember Mushrush, Amber N. Coats, Sara Dao, Grace Dittmar, Taylor Gore, Taiya M. Jarva, Giorgi Kenkebashvili, Sudiksha Rathan-Kumar, Gabriella M. Reyes, Garrett L. Watts, Victoria Kalene Watts, Deena Dubrow, Gabrielle Lewis, Benjamin H. Stone, Bingjie Xue, Steven G. Cresawn, Dmitri Mavrodi, Viknesh Sivanathan, Danielle Heller

## Abstract

During infection, bacteriophages produce diverse gene products to overcome bacterial anti-phage defenses, to outcompete other phages, and take over cellular processes. Even in the best-studied model phages, the roles of most phage-encoded gene products are unknown, and the phage population represents a largely untapped reservoir of novel gene functions. Considering the sheer size of this population, experimental screening methods are needed to sort through the enormous collection of available sequences and identify gene products that can modulate bacterial behavior for downstream functional characterization. Here, we describe the construction of a plasmid-based overexpression library of 94 genes encoded by Hammy, a Cluster K mycobacteriophage closely related to those infecting clinically important mycobacteria. The arrayed library was systematically screened in a plate-based cytotoxicity assay, identifying a diverse set of 24 gene products (representing 25% of the Hammy genome) capable of inhibiting growth of the host bacterium *Mycobacterium smegmatis*. Half of these are related to growth inhibitors previously identified in related phage Waterfoul, supporting their functional conservation; the other genes represent novel additions to the list of known anti-mycobacterial growth inhibitors. This work, conducted as part of the HHMI-supported Science Education Alliance Gene-function Elucidation by a Network of Emerging Scientists (SEA-GENES) project, highlights the value of parallel, comprehensive overexpression screens in exploring genome-wide patterns of phage gene function and novel interactions between phages and their hosts.

## Introduction

Bacteriophages and bacteria are locked in an ancient, high-stakes arms race, giving rise to myriad genetic strategies for infecting or thwarting infection, respectively (Hampton *et al*. 2020). Phage genomes encode a core set of gene products that form the virion structure or interact with host factors to carry out key steps of infection, including genome replication, gene expression, and cell envelope disruption. Typically, phage also encode a suite of highly variable accessory genes thought to confer an advantage under specific conditions (Dedrick *et al*. 2013, 2017; Hatfull 2020). Comparative analyses of many sequenced phages have revealed a continuum of genetic diversity, with mosaic genomes encoding thousands of different phamilies, or groupings of related gene products sharing >32% amino acid identity; most phamilies have no recognizable sequence features and cannot be assigned function (Cresawn *et al*. 2011; Pope *et al*. 2015, 2017; Hatfull 2020; Gauthier *et al*. 2022). In turn, bacteria have evolved a multitude of systems that sense and respond to phage infection via either abortive or non-abortive mechanisms (Hampton *et al*. 2020; LeRoux and Laub 2022; Millman *et al*. 2022). Recent work has revealed that several of these abortive systems are specifically triggered by core phage products including capsid structures (Huiting *et al*.; Tal *et al*. 2021; Zhang *et al*. 2022) and DNA replication machinery (Stokar-Avihail *et al*. 2023). Phages are also in vicious competition with each other, and increasingly, many phage products are thought to interact with host factors to block infection by competitors (Dedrick *et al*. 2017; Ko and Hatfull 2018). It is evident that the interactions between phages and bacteria are complex, and further work is needed to elucidate the full expanse of genetic functions that both sides have evolved.

One powerful strategy for investigating phage gene function is overexpression in the bacterial host, connecting genes to phenotypes like inhibition of bacterial growth and serving as a first experimental filter to identify phage products that interact with key host factors (Liu *et al*. 2004; Molshanski-Mor *et al*. 2014; Ko and Hatfull 2018). Recent overexpression screens have shown that genes capable of disrupting mycobacterial growth are abundant within the mycobacteriophage population (Ko and Hatfull 2020; Heller *et al*. 2022). Systematic overexpression of each gene encoded by Cluster K mycobacteriophage Waterfoul revealed that roughly one-third of gene products hindered *Mycobacterium smegmatis* growth (Heller *et al*. 2022). Given the remarkable size and diversity of the phage population, additional phenotypic mining of phage genomes is sure to uncover many more genes capable of modulating bacterial behavior.

Here, we report the results of a genome-wide overexpression screen for Hammy, a temperate siphovirus that infects *M. smegmatis*. Based on gene content similarity, Hammy is classified as part of Cluster K (Subcluster K6) and is related to Waterfoul and several clinically relevant mycobacteriophages such as ZoeJ (Subcluster K2) (Russell and Hatfull 2016; Anders *et al*. 2017; Dedrick *et al*. 2019b; Guerrero-Bustamante *et al*. 2021). The Hammy genome contains 95 predicted protein-coding genes, only ∼35% of which have annotated functions (Figure 1) (Anders *et al*. 2017). Out of 94 Hammy genes evaluated in our systematic screen, a diverse set of 24 genes, representing 25% of the genome, were found to interfere with host growth; 12 of these diverse products caused near complete growth abolition. This study, done as part of the Science Education Alliance Gene-function Exploration by a Network of Emerging Scientists, or SEA-GENES project (Heller and Sivanathan 2022), contributes novel hits to the growing list of known anti-mycobacterial proteins while also providing insight into the conservation of host inhibition across related phage genomes.

**Figure 1:**
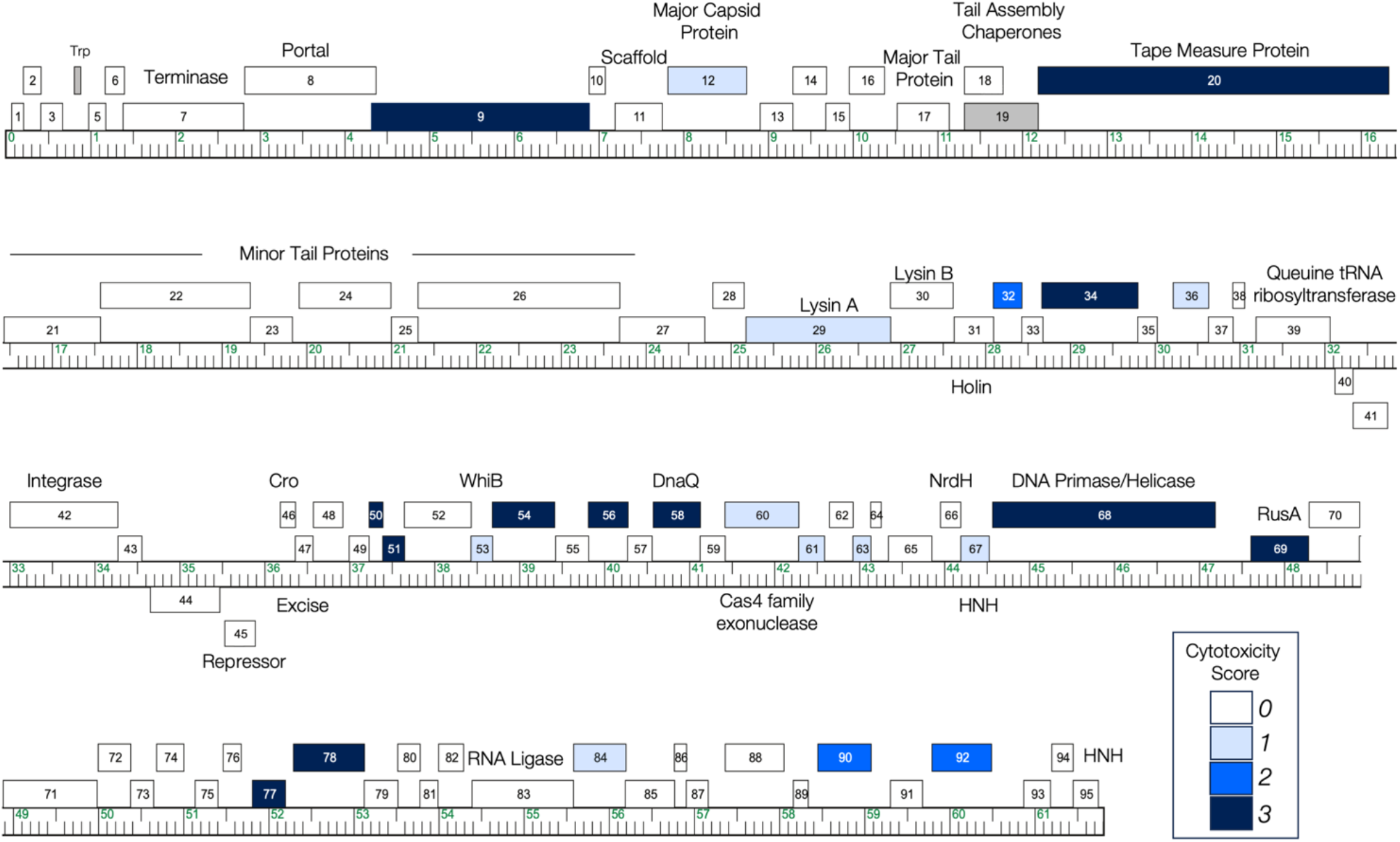
The genome of phage Hammy. The Hammy genome is shown as a ruler with kbp markers and genes represented by boxes—those above the line are transcribed rightwards and those below are transcribed leftwards. Numbers inside the box correspond to gene numbers and predicted functions are indicated above each gene. Box shading corresponds to cytotoxicity scoring, with white boxes designating genes found to have no effect on *M. smegmatis* growth (cytotoxicity score *0*), hatched box indicating omitted gene *19* not tested in this study, and blue representing observed toxicity in our assay. The saturation of blue boxes corresponds to the severity of growth inhibition using the following scores: light blue (score *1*; reduction in colony size; genes *12, 29, 36, 53, 60, 61, 63, 67*, and *84*), medium blue (score *2*; 1–3 log reduction in viability; genes *32, 90*, and *92*), and dark blue (score *3*; >3-log reduction in viability; genes *9, 20, 34, 50, 51, 54, 56, 58, 68, 69, 77*, and *78*).

## Materials and Methods

### Growth of mycobacteria and mycobacteriophage

*M. smegmatis* mc^2^155 was grown at 37 °C in Middlebrook 7H9 (Difco) broth supplemented with 10% AD (2% w/v Dextrose, 145 mM NaCl, 5% w/v Albumin Fraction V), 0.05% Tween80, and 10 µg/ml cycloheximide (CHX) or on Middlebrook 7H10 (Difco) or 7H11 (Remel) agar supplemented with 10% AD and 10 µg/ml CHX. For transformation of *M. smegmatis* mc^2^155, electrocompetent cells were electroporated with ∼50-100 ng of pExTra plasmid DNA, recovered in 7H9 broth for 2 h at 37 °C with shaking, and transformants selected on 7H10 or 7H11 agar supplemented with 10 µg/ml Kanamycin (GoldBio). After 4 days of incubation at 37 °C, colonies of transformants were used directly in plate-based cytotoxicity assays or to inoculate cultures for liquid growth assays. Hammy was propagated on *M. smegmatis* mc^2^155 grown at 25 °C in the presence of 1 mM CaCl_2_ and no Tween in Middlebrook media and top agar.

### Construction of the pExTra Hammy Library

Each Hammy gene was cloned into the pExTra shuttle vector (Heller *et al*. 2022) downstream of an anhydrotetracycline-inducible promoter, *pTet* (Ehrt *et al*. 2005), to control gene expression, and upstream of linked *mcherry* transcriptional reporter (Figure 2A). Genes were PCR-amplified (New England Biolabs Q5 HotStart 2X Master Mix) from a high-titer Hammy lysate using a forward primer complementary to the first 15-25 bp of each gene sequence (Integrated DNA Technologies), introducing a uniform ATG start codon, and a reverse primer complementary to the last 15-25 bp of the gene sequence, including a uniform TGA stop codon (Supplemental Table 1). All forward primers contained a uniform, RBS-containing 5**’** 21 bp sequence and all reverse primers contained a separate 5’ 25 bp sequence; these added sequences have identity to the pExTra plasmid flanking the site of insertion. Linearized pExTra plasmid was prepared via PCR (NEB Q5 HotStart 2X Master Mix) of pExTra01 (Heller *et al*. 2022) using divergent primers pExTra_F and pExTra_R and assembled with each gene insert by isothermal assembly (NEB HiFi 2X Master Mix).

**Figure 2:**
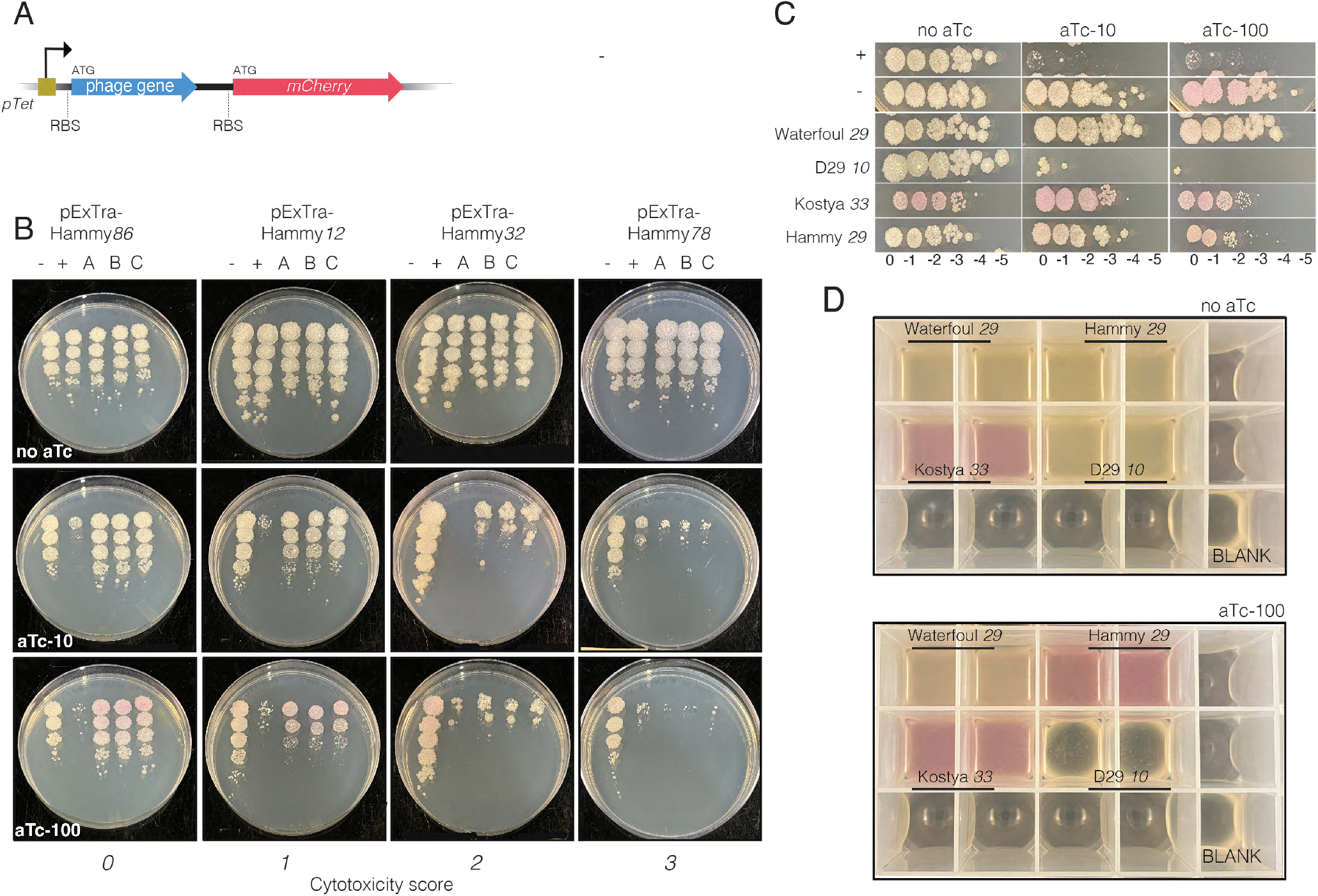
Expression of phage genes from the pExTra plasmid. A) Recombinant pExTra plasmids constructed in this study encode Hammy gene sequences downstream of the *pTet* promoter and upstream of *mcherry*. The two genes in this pExTra operon are transcriptionally linked, each bearing their own translational signals. B) Results of representative cytotoxicity assays are shown to demonstrate the range of observed growth defects. In each assay, colonies of *M. smegmatis* mc^2^155 transformed with the specified pExTra plasmid were resuspended, serially diluted, and spotted on 7H11 Kan media containing 0, 10, or 100 ng/ml aTc. Triplicate colonies were tested for each gene alongside a positive control strain (+) transformed with pExTra02 (expressing wildtype Fruitloop *52*) and a negative control strain (-) transformed with pExTra03 (expressing Fruitloop *52 I70S*). *lysA* genes from various mycobacteriophages were expressed from pExTra and tested C) in the plate-based cytotoxicity assay alongside pExTra02 (+) and pExTra03 (-) controls and D) in an end-point liquid growth assay. Two colonies of each transformed *M. smegmatis* strain were grown until saturation and sub-cultured in duplicate 24-well plates. Upon reaching an OD600 of ∼0.2, one set of cultures was induced by addition of 100 ng/ml aTc and the other left as an uninduced control. Growth was monitored over 24 h of induction at 37 °C with shaking. Plate images from a representative experiment showing lysis and culture appearance after 24 h induction are shown.

D29 *10* and codon-optimized Kostya *33* inserts were synthesized with 5’ and 3’ flanking sequences (Integrated DNA Technologies) and assembled directly with linearized pExTra plasmid. Recombinant plasmids were recovered by transformation of *E. coli NEB5α F’I*^*Q*^ (New England Biolabs) and selection on LB agar supplemented with 50 µg/ml Kanamycin. The pExTra-Waterfoul29 plasmid was described previously (Heller *et al*. 2022).

The inserted genes for all recovered pExTra plasmids were sequence-verified by Sanger sequencing (Azenta) using sequencing primers pExTra_universalR and pExTra_seqF; longer genes were also sequenced with internal sequencing primers listed in Supplemental Table 1. All plasmid inserts were found to match the published genome sequence.

### Cytotoxicity Screening and Phenotype Scoring

To assess cytotoxicity, pExTra-transformed colonies were resuspended and serially diluted in 7H9 broth then spotted on 7H10 or 7H11 plates supplemented with 10 µg/ml Kanamycin and 0, 10, or 100 ng/ml anhydrotetracycline (aTc; Alfa Aesar). Each strain was tested in triplicate alongside the pExTra02 positive control plasmid, encoding cytotoxic gene Fruitloop *52*, and the pExTra03 negative control plasmid, encoding a non-toxic mutant allele of Fruitloop *52* (I70S) (Ko and Hatfull 2018; Heller *et al*. 2022). Growth was monitored over 5 days at 37 °C; typically, spot growth was visible after 2 days at 37 °C with effects on colony size more apparent after 3-5 days of incubation. Cytotoxic phenotypes were scored by comparing the spot dilution out to which cells grew in the presence versus the absence of aTc inducer and classified as either having no effect (score 0), being moderately cytotoxic with a 1-3 log reduction in cell viability (score 2), or being highly cytotoxic, causing complete or near complete (>3-log) inhibition of growth (score 3). Strains were also evaluated for aTc-dependent size reduction in individual colonies (score 1) as compared to the Fruitloop *52-I70S* negative control strain on the same aTc plate and the same strain on plates without inducer. Pink colony color from *mcherry* gene expression provided a visual indicator of gene expression through the *pTet* operon.

All genes were tested in at least two independent cytotoxicity experiments; one round of screening was done on 7H10 agar and another on 7H11 agar, with good agreement observed between the different growth media. Unless otherwise indicated, the results and images reported in this study are all from experiments done on 7H11 agar. Reported cytotoxic genes were found to cause growth inhibition in more than two independent experiments with strong agreement between triplicate samples within each experiment. The magnitude of cytotoxicity for some genes was observed to vary slightly between experiments, likely due to minor variations in media or growth conditions (e.g., compare Hammy *12* results in Figure 2B and Supplemental Figure 1); variability was more pronounced in those strains observed to have milder toxic effects, which are potentially at the detection limit of our assay. If differences in magnitude were observed between experiments, genes were scored based on the more conservative result. The inclusion of the Fruitloop *52* control strains on each screening plate aided in the evaluation of relative gene-mediated effects, and results were ultimately scored based on observations on 100 ng/ml aTc.

For liquid growth assays, transformants were grown in Middlebrook 7H9 broth with 10 µg/ml Kanamycin until saturated. Cultures were back-diluted in fresh medium and induced with 100 ng/ml aTc at an O.D. 600 of ∼0.2-0.4; growth wasmonitored at 37 °C with shaking.

### Hammy genomic analysis

The Hammy genome map was created using the web-based tool Phamerator (phamerator.org). Reported gene functions are based on those available in the Hammy GenBank record (Accession KY087993). In several cases, these GenBank annotations were supplemented by comparison of functional assignments for other phamily members on PhagesDB (Russell and Hatfull 2016), especially those from more recently annotated Cluster K mycobacteriophage genomes; these designations were confirmed using HHPRED (PDB_mmCIF70, SCOPe70_2.08, Pfam-A_v35, NCBI_Conserved_Domains(CD)_v3.19) (Gabler *et al*. 2020), NPS Helix-Turn-Helix predictor (https://npsa-prabi.ibcp.fr/), and DeepTMHMM (Hallgren *et al*. 2022). Gene content comparison between Hammy and Waterfoul genomes was performed using the gene content comparison tool on phagesDB (https://phagesdb.org/genecontent/), with phamily designations downloaded from the database on February 27, 2023. Reported gene product similarities were determined by multiple sequence alignment with Clustal Omega (Madeira *et al*. 2022).

## Results and Discussion

### A system for studying Hammy gene overexpression in *M. smegmatis*

To systematically investigate the effects of Hammy gene overexpression on mycobacterial growth, an arrayed library of Hammy protein-coding genes was generated. Ninety-four genes were included in this analysis, excluding only gene *19*, the long isoform of the tail assembly chaperone produced through programmed frameshifting; this analysis also excluded the single tRNA encoded in the Hammy genome. Each gene was cloned into the multi-copy expression vector, pExTra, under the control of aTc-inducible promoter *pTet* and transcriptionally linked to fluorescent reporter gene *mcherry* (Figure 2A) (Heller *et al*. 2022). Sequence-verified plasmids were used to transform *M. smegmatis* mc^2^155, and transformants for all 94 strains were analyzed in a semi-quantitative spot dilution assay to measure the impacts of overexpressing each gene on bacterial growth.

As illustrated in Figure 2B, for each Hammy gene, dilution series for three transformed colonies were prepared and spotted on increasing concentrations of aTc inducer alongside control cells expressing a wildtype (pExTra02) or mutant (pExTra03) allele of phage Fruitloop gene *52*. Previous work has established that overproduction of wildtype Fruitloop gp52 severely impairs growth of *M. smegmatis*, whereas comparable levels of the gp52-I70S variant cause no reduction in growth (Ko and Hatfull 2018). An aTc-dependent increase in pink coloration of spots was observed for the negative control pExTra03 strain and several experimental strains, confirming previous reports of tunable expression from *pTet* (Ehrt *et al*. 2005; Parikh *et al*. 2013; Heller *et al*. 2022). For Hammy genes *46* and *71*, aTc-independent *mcherry* expression was observed (Supplemental Figure 1), suggesting that these sequences may harbor promoter sequences that can be recognized by the host transcriptional machinery.

Our screening strategy allowed for observation of a range of impacts on host growth, from mild reduction in colony size to multi-log reduction in colony number, and growth effects were scored using an index from *0* (no effect on bacterial growth) to *3* (near complete abolition of host growth) (see Figure 2B for representative examples). In our system, cytotoxicity was observed to increase for some genes in a dose-dependent manner on 10 and 100 ng/ml aTc (e.g., Hammy *29*; Supplemental Fig 1), whereas for others, full cytotoxic effects were seen at the lower aTc concentration (e.g., Hammy *67*; Supplemental Figure 1). Final cytotoxicity scores were assigned based on observations from the higher concentration of inducer.

### Overexpression of most Hammy genes does not inhibit host growth

Of the 94 Hammy genes screened, 74% caused no appreciable reduction in bacterial growth in our assay (cytotoxicity score *0*), with comparable viability and colony size in the presence or absence of aTc inducer (Figure 2B, Supplemental Figure 1). This figure is in line with a previous systematic screen of genes encoded by related mycobacteriophage Waterfoul, 66% of which had no measurable impact on *M. smegmatis* growth (Heller *et al*. 2022), as well as a previous study by Ko and Hatfull that found that roughly 75% of a collection of genes with no known function (NKF) from assorted mycobacteriophages were not toxic when overexpressed in *M. smegmatis* (Ko and Hatfull 2020). Out of the 70 Hammy genes found to be non-toxic in our assay, 60 displayed varying degrees of pink spot coloration on the highest concentration of inducer (Supplemental Figure 1). This visual confirmation of transcription and translation occurring through the *pTet* operon suggests that phage gene expression is occurring in most strains; however, we cannot rule out that some of these gene products may not accumulate within cells in our assay.

### A diverse set of Hammy gene products inhibit *M. smegmatis* growth

Out of 94 Hammy protein-coding sequences, 24 genes were found to reproducibly reduce growth of *M. smegmatis* in an aTc-dependent manner (Table 1, Figure 1, Supplemental Figure 1). Within this set, 9 genes caused effects characterized as mild, with notable reduction in colony size but no measurable decrease in plating (cytotoxicity score *1*). The remaining 15 genes caused severe growth defects, with 3 causing a 1-2 log reduction in colony number (cytotoxicity score *2*), and 12 resulting in >3-log reduction in colony number or almost complete inhibition of growth (cytotoxicity score *3*). These 24 cytotoxic gene products range in size from 84 amino acids to the largest gene product encoded in the genome, the 4,140 amino acid tape measure protein gp20 (Table 1).

**Table 1:** Hammy genes observed to inhibit *Mycobacterium smegmatis* growth upon overexpression.

| Gene | Phamily<sup>1</sup> | Length (aa) | Predicted function and features | Cytotoxicity<sup>2</sup> |
| --- | --- | --- | --- | --- |
| 12 | 68545 | 930 | Major capsid protein | 1 |
| 29 | 68638 | 1704 | Lysin A | 1 |
| 36 | 68706 | 420 | NKF | 1 |
| 53 | 68569 | 84 | WhiB family transcription factor | 1 |
| 60 | 68645 | 289 | Cas4 family exonuclease | 1 |
| 61 | 3651 | 101 | NKF | 1 |
| 63 | 67008 | 213 | NKF | 1 |
| 67 | 60651 | 336 | HNH endonuclease | 1 |
| 84 | 69022 | 204 | NKF | 1 |
| 32 | 67117 | 339 | NKF; TMD<sup>3</sup> | 2 |
| 90 | 14302 | 627 | NKF | 2 |
| 92 | 59346 | 699 | NKF | 2 |
| 9 | 68668 | 2574 | NKF | 3 |
| 20 | 67034 | 4140 | Tape measure protein; TMD | 3 |
| 34 | 65534 | 1134 | MRE11 double-strand break endo/exonuclease | 3 |
| 50 | 44663 | 165 | NKF; TMD | 3 |
| 51 | 850 | 258 | NKF | 3 |
| 54 | 68730 | 735 | NKF; DUF3310 | 3 |
| 56 | 68998 | 468 | NKF | 3 |
| 58 | 67001 | 555 | DnaQ-like (DNA polymerase III subunit) | 3 |
| 68 | 607 | 2622 | DNA primase/helicase | 3 |
| 69 | 749 | 678 | RusA-like resolvase | 3 |
| 77 | 3303 | 390 | NKF | 3 |
| 78 | 596 | 834 | NKF; TMD; HTH-DNA-binding | 3 |

<sup>1</sup>Phamily numbers were recorded from phagesdb.org as of February 27, 2023.
<sup>2</sup> Growth of strains on media supplemented with 10 ng/ml or 100 ng/ml aTc was compared to the same strains plated on media without aTc. Those scored as 0, had no effect on host growth at aTc-10; those scored as 1 exhibit an aTc-dependent reduction in colony size, those denoted as 2 demonstrate a ~1-3-log difference in growth in the presence of aTc, and those denoted as 3 demonstrate a severe >3-log reduction in growth in the presence of aTc.
<sup>3</sup> TMD= Transmembrane domain

Only a subset of these gene products have putative functional assignments or recognizable sequence features (Table 1). Two are structural proteins, with overproduction of the major capsid subunit gp12 causing mild toxicity (score *1*) and overproduction of the tape measure protein gp20 causing a severe growth defect (score *3*).

Interestingly, overproduction of the Lysin A protein from Hammy (gp29) was found to cause a mild growth defect (Figure 2C; Supplemental Figure 1). Efficient phage-mediated lysis of the mycobacterial cell is thought to require coordination between the phage holin, peptidoglycan-hydrolyzing Lysin A, and Lysin B esterase protein to disrupt the multi-layered cell envelope (Catalão and Pimentel 2018). Expression of *lysA* genes by themselves is typically not toxic (Payne *et al*. 2009; Payne and Hatfull 2012; Catalão and Pimentel 2018; Heller *et al*. 2022); however, a small number of mycobacteriophage *lysA* gene phamilies have been reported to cause varying degrees of host cell death in a holin-independent manner when independently overexpressed from plasmids (Payne and Hatfull 2012).

Six gene products with predicted functions related to DNA metabolism were observed to hinder host growth, including the predicted MRE11 double-strand break exo/endonuclease (gp34), Cas4-family exonuclease (gp60), RusA-like resolvase (gp69), one of two Hammy HNH endonucleases (gp67), and candidate replisome components DNA primase/helicase (gp68) and DnaQ (gp58). Several of these were among the most toxic gene products scored in our assay, with overproduction of the MRE11 double-strand break exo/endonuclease (gp34), DnaQ (gp58), DNA primase/helicase (gp68), and RusA (gp69) causing near complete abolition of host growth (Table 1, Supplemental Figure 1). Another highly cytotoxic gene product, NKF protein Hammy gp54, contains a conserved domain of unknown function (DUF3310), which was previously implicated in nucleotide kinase activity for the T7 coliphage protein gp1.7 (Tran *et al*. 2008, 2012, 2014).

Three genes with putative DNA-binding functions were observed to inhibit host growth (Table 1). This includes the WhiB transcription factor gp53 and predicted helix-turn-helix DNA-binding protein, gp36, both of which caused a reduction in colony size upon overproduction (score *1*). The potent growth inhibitor gp78 (score *3*), is predicted to harbor a C-terminal helix-turn-helix DNA-binding domain as well as 4 putative transmembrane helices towards the N-terminus of the protein. Two additional gene products (gp32 and gp50) are predicted by DeepTMHMM to contain transmembrane domains (TMD). Hammy gp32 contains a single predicted TMD at its N-terminus and is encoded immediately downstream of the candidate holin protein, gp31, which has four predicted TMDs. Phage lysis cassettes often encode multiple holin-like TMD proteins (Hatfull 2018; Pollenz *et al*. 2022), and gp32 may thus play a role in disruption of the cell envelope. The small, 54-amino acid protein gp50 is predicted to have a 20-amino acid TMD at its C-terminus such that a short cytoplasmic domain at the N-terminus would be anchored in the cell membrane. Predicted membrane proteins, notorious for impairing cell growth when produced at high levels in bacteria (Miroux and Walker 1996; Wagner *et al*. 2006), were not uniformly toxic in our cytotoxicity screen. In total, Hammy encodes 7 proteins with predicted TMDs. Overproduction of three of these (gp31, gp35, gp44) had no detectable effect on host growth in our assay, though we note that obvious pink coloration of spots was only observed for one of these (gp44; Supplemental Figure 1).

Overall, our systematic screen revealed that one-quarter of Hammy genes are deleterious to *M. smegmatis* growth when overexpressed in our system, representing a set of mycobacterial growth inhibitors that are diverse in terms of their size, sequence features, and magnitude of effect. As with any overexpression screen, it is possible that some of the observed growth defects, especially those milder defects seen only at the higher level of induction, may be a consequence of artificially high protein levels. It is encouraging though that half of the hits described here caused near complete abolition of host growth, with severe growth defects observed even on the lower concentration of aTc (Supplemental Figure 1). These products offer promising candidates for further elucidation of the interactions between phage factors and key cellular complexes.

### Conservation of mycobacterial growth inhibition by related phage genes

To facilitate gene content comparisons across highly mosaic phage genomes, gene products encoded within sequenced actinobacteriophage genomes are grouped into gene phamilies based on amino acid similarity (Cresawn *et al*. 2011), and each gene in the Hammy genome belongs to a designated phamily with known homologues in other phages (Russell and Hatfull 2016). A systematic screen using the same pExTra overexpression system, plate-based cytotoxicity assay, and scoring index was recently reported for related Cluster K mycobacteriophage Waterfoul (Heller *et al*. 2022), enabling a comparison of phenotypes conferred by shared gene phamilies encoded by the two related phages. Waterfoul and Hammy encode 46 phamilies in common (Cresawn *et al*. 2011; Russell and Hatfull 2016), and 34 of these were consistently classified as toxic or not toxic in this study and the Waterfoul report (Figure 3). Nine shared gene phamilies were observed to inhibit mycobacterial growth in both studies, including three highly cytotoxic phamilies encoded in both genomes represented by Hammy NKF genes *9, 54*, and *78* (Figure 3). Consistent cytotoxic effects by homologous gene products further support these phamilies as strong candidates for factors targeting essential mycobacterial processes, and in general, functional comparisons offer opportunities to probe the conserved sequence determinants of host interference.

**Figure 3:**
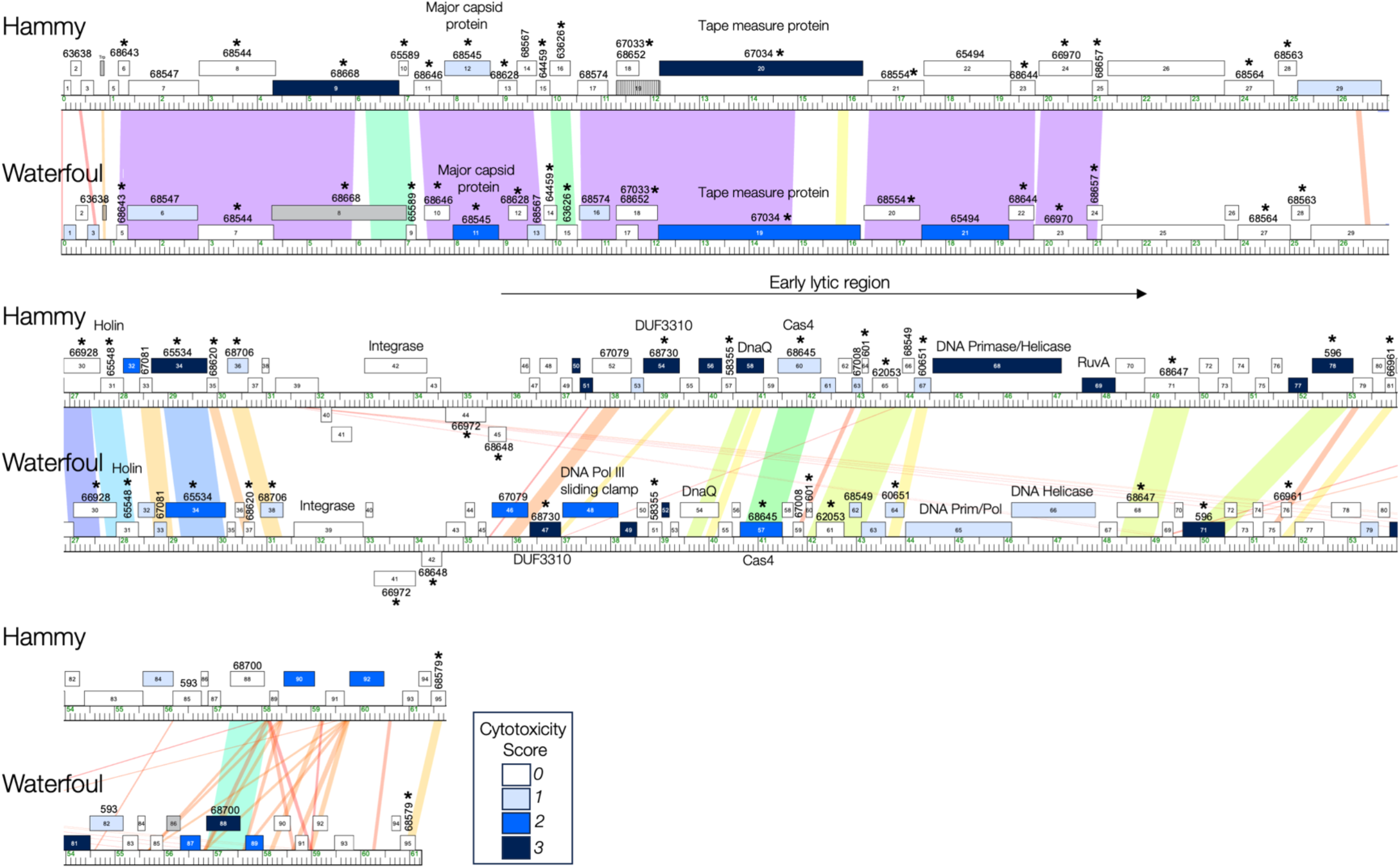
Comparison of Hammy and Waterfoul phenotypes. Aligned maps of the Hammy and Waterfoul genomes are shown, with gene shading corresponding to the cytotoxicity scores for Hammy genes as reported in Figure 1 of this report and Waterfoul data reported by Heller et al. (Heller *et al*. 2022), with pairwise nucleotide sequence similarity shown as spectrum-colored shading between genomes. Violet indicates the highest level of similarity, and red the least similarity, above a threshold E-value of 10^−4^. Gene phamilies found in both genomes are labeled by phamily numbers (taken from phagesDB.org as of February 27, 2023) with asterisks indicating those gene phamilies that were found to be cytotoxic (scores *1-3*) or non-toxic consistently across both genomes.

Most of the phenotypic discrepancies between shared Hammy and Waterfoul phamilies are for genes causing only reduction in colony size (score *1*) in one of the two reports (Figure 3), suggesting that these milder growth defects may be at the detection limit of our assay. However, a more notable discrepancy in phenotype was observed for 3 shared gene phamilies: Hammy genes *22, 52*, and *88* were found to have no impact on host growth when overexpressed, whereas the homologous genes from Waterfoul (genes *21, 46*, and *88*) caused severe reduction in colony number (score 2 or 3) upon overexpression (Figure 3). It remains to be seen whether these discrepancies are due to variation in biological function or are a result of expression differences in our system. For example, major tail protein Hammy gp21 is 78% identical to cytotoxic protein Waterfoul gp22, but spots overproducing Hammy gp21 displayed no pink coloration on aTc, suggesting that the lack of toxic phenotype may be due to insufficient protein levels (Supplemental Figure 1).

Interestingly, consistent toxicity was observed for two conserved structural components—the related major capsid proteins Hammy gp11 and Waterfoul gp12 (80% identical) and TMD-containing tape measure proteins Hammy gp20 and Waterfoul gp19 (58% identical). Virion structures are typically thought to function outside or at the surface of the cell making their cytoplasmic cytotoxicity a somewhat surprising result; however, these structural components are highly expressed in the cell during lytic infection (Dedrick *et al*. 2013, 2019a), and recent work has demonstrated their potential to trigger diverse cellular responses (Huiting *et al*.; Pedulla *et al*. 2003; Zhang *et al*. 2022; Stokar-Avihail *et al*. 2023). Thus, further examination of the cytotoxicity conferred by these structural genes as well as impacts caused by other mycobacteriophage structures is warranted.

The mild impact on growth caused by WhiB protein Hammy gp53 is consistent with a previous finding that related phamily member, TM4 gp49 (45% identical), disrupts cell division and inhibits growth when overproduced in *M. smegmatis* (Rybniker *et al*. 2010). Rybniker et al. proposed that the TM4 WhiB impaired growth by downregulating expression of the essential host transcription factor *whiB2* through competitive binding of the *whiB2* promoter (Rybniker *et al*. 2010), and the Hammy WhiB may hinder host growth through a similar mechanism.

Phenotypic comparisons can also be made between genes that belong to distinct gene phamilies but are predicted to carry out common functions. Different phamilies with putative roles in Hammy or Waterfoul genome replication were identified as cytotoxic, including the Waterfoul DNA primase/polymerase gp65, the Waterfoul DNA helicase protein gp66, the Hammy DNA primase/helicase gp68, as well as predicted sliding clamp subunit Waterfoul gp48, and DnaQ protein Hammy gp58 (Figure 3). We note that the Waterfoul DnaQ protein gp54, which is categorized in a separate phamily from Hammy DnaQ gp58, was not observed to be toxic, and thus, cytotoxicity may be specific to the Hammy gp58 phamily rather than a general consequence of overproduction of phage DnaQ proteins. Here too, additional investigation is needed to understand whether cytotoxicity by phage replication proteins is due to interference with the host DNA or replication machinery or because of some other mechanism. It is worth noting that in recent explorations of phage-host dynamics, phage-encoded replication proteins have also been observed to elicit diverse cellular defense responses (Stokar-Avihail *et al*. 2023).

Hammy and Waterfoul vary in their lysis cassettes, with divergent *lysA* genes adjacent to *lysB* and holin-encoding genes from common gene phamilies (Figure 3). Overproduction of related Lysin B (75% identical) and holin (78% identical) proteins from Wateroul and Hammy had little to no effect on host growth, whereas the distinct *lysA* genes had differing impacts on host growth; Hammy *lysA* overexpression was mildly toxic, causing a reduction in colony size, whereas Waterfoul *lysA* overexpression had no visible impact on host growth. Most Lysin A proteins tested to date, including the Hammy Lysin A homolog in phage Bxz2 (48% amino acid identity) do not cause cell lysis independent of holin activity; however, a small number of mycobacteriophage *lysA* genes have previously been shown to cause holin-independent lysis (Payne and Hatfull 2012). Using an *M. smegmatis* liquid growth and ATP release assay, Payne and Hatfull reported that overproduction of the D29 Lysin A protein gp10 caused pronounced ATP release and cell lysis, whereas overproduction of other Lysin A proteins from other gene phamilies, including Waterfoul gp29 homolog, Kostya gp33, had only mild effects.

To directly compare the impacts of Hammy *lysA* overexpression on *M. smegmatis* growth to those of previously studied mycobacteriophage *lysA* genes, we tested several *lysA* genes in both our plate-based cytotoxicity assay and a liquid end-point growth assay. Consistent with the holin-independent lysis results reported by Payne and Hatfull, D29 *lysA* overexpression completely inhibited growth in both assays (Figure 2C and 2D), with signs of cell lysis, including cleared culture and visible cell debris, observed in liquid after 24 h induction. The effect mediated by D29 *lysA* expression was much more pronounced than that observed for the Hammy or Kostya *lysA* genes, both of which only caused an aTc-dependent reduction in colony size on plates (Figure 2C) but no notable signs of cell death in liquid (Figure 2D). Pink coloration of cultures expressing the Hammy *lysA* in the presence of inducer suggests that expression through the *pTet* operon is occurring in cells. Interestingly, even in the absence of aTc, cells transformed with the pExTra-Kostya33 plasmid turned pink and formed smaller colonies, suggesting that this *lysA* sequence may harbor an internal gene start (Catalão *et al*. 2011). As seen previously, despite being 66% identical to the Kostya Lysin A protein, overproduction of the Waterfoul Lysin A had no effect on host growth on plates (Figure 2C) or in liquid (Figure 2D), though only faint pink color was observed in the presence of aTc, meaning that the Waterfoul *lysA* may be poorly expressed in our system. These data imply that although mildly toxic when produced independently on plates, the Hammy Lysin A is not mediating pronounced holin-independent cell lysis in our system like is seen with the D29 Lysin A. Mycobacteriophage Lysin A proteins are multi-domain proteins, with putative peptidase, glycoside hydrolase, and cell wall-binding functions (Payne and Hatfull 2012). The additional observation of holin-independent cytotoxicity presented here may prove useful in genetically dissecting the determinants of these lytic and non-lytic phenotypes.

Finally, 12 of the 24 growth inhibitors identified in our Hammy screen (Hammy *32, 50, 51, 56, 58, 61, 68, 69, 77, 84, 90, 92*) represent gene phamilies for which, to the best of our knowledge, cytotoxic effects have not been previously reported, thus also adding many novel hits to the set of known phage-encoded growth inhibitors.

### Insights into genomic patterns of mycobacterial growth inhibition

Beyond phenotypic comparisons of shared gene phamilies, our systematic Hammy screen results also lend support for several genome-wide patterns first observed for Waterfoul (Heller *et al*. 2022). Cytotoxic genes are abundant within phage genomes: one-quarter of Hammy genes and one-third of Waterfoul genes were found to inhibit growth of the host when overexpressed, with a similar fraction of genes in each genome capable of causing severe growth defects in the host (Figure 3). Both NKF genes and genes with predicted functions inhibit host growth, and while it is unlikely that all are employed during infection to kill the host cell, this high frequency of host interference underscores the complexities of host-phage dynamics. Indeed, recent work on the diverse interactions that occur between phages infecting a common host, suggests that some of these cytotoxic phage products may bind cellular factors with the purpose of blocking secondary infection by other phages (Dedrick *et al*. 2017; Ko and Hatfull 2018; Gentile *et al*. 2019; Montgomery *et al*. 2019).

We also find that cytotoxic genes are distributed throughout both genomes and are often found near other cytotoxic genes. Transcriptomic data from related Cluster K mycobacteriophage ZoeJ suggests that many of these cytotoxic genes are likely co-expressed over the course of the Hammy or Waterfoul life cycle (Dedrick *et al*. 2019a), and it is plausible that cytotoxic products encoded by adjacent genes target related host processes. The predicted early lytic regions of both genomes are enriched with cytotoxic genes, including conserved DUF3310 genes as well as various genes predicted to function in transcription, DNA replication, and DNA metabolism (Figure 3). As some of the first gene products made during infection, these are good candidates for factors involved in the coordinated takeover of critical cellular processes. A second cytotoxic gene cluster is found between the holin and integrase genes in both Hammy and Waterfoul, with cytotoxic genes encoding poorly conserved TMD proteins Hammy and Waterfoul gp32 (19% amino acid identity) flanked by conserved holin genes and cytotoxic nucleases, Hammy and Waterfoul gp34. Phage lysis cassettes are highly mosaic (Hatfull 2018; Pollenz *et al*. 2022), and phenotypic profiling such as that described here may identify convergent functions within these cassettes and help to better characterize the mycobacteriophage lysis pathway.

In conclusion, only a small fraction of the thousands of known mycobacteriophage gene phamilies have been experimentally explored to date (Rybniker *et al*. 2008, 2010, 2011; Mehla *et al*. 2017; Ko and Hatfull 2018, 2020; Heller *et al*. 2022). Here, we show that systematic screening of phage genomes in a simple, plate-based overexpression assay can identify a diverse set of bacterial effectors, offering many promising candidates for elucidation of novel interactions between phages and their mycobacterial hosts. Extension of this screening strategy to additional diverse mycobacteriophage genomes will undoubtedly uncover many more novel effectors and provide additional insight into the patterns of phenotypic conservation examined here.

## Data Availability

All plasmids and plasmid sequences reported in this study are available upon request. The authors affirm that all data necessary for confirming the conclusions of this article are represented fully within the article and its tables and figures. Extended data can be found at genesDB.org.

## Acknowledgements

This work was conducted as part of the HHMI-supported Science Education Alliance GENES (Gene-function Exploration by a Network of Emerging Scientists) project. We would like to acknowledge several members of the Science Education Alliance for their contributions to this work, including Billy Biederman, Debbie Jacobs-Sera, Dan Russell, J.C. Gardner III, Kathleen L. Johnson, Jasmine J. Ransom, Chazmyn R. Riley, Daniel T. Sinclair, Breanna S. Smith, Audra E. Thompson, Savannah L. Underwood, and all faculty and student members of the SEA-GENES program. We would like to thank Graham Hatfull for reagents, research advice, and helpful input on the manuscript. We thank New England Biolabs (NEB) and Integrated DNA Technologies (IDT) for providing reagent support.

