## Supplemental Materials for "A genome-wide overexpression screen reveals *Mycobacterium smegmatis* growth inhibitors encoded by mycobacteriophage Hammy"

**Supplemental Figure 1: Systematic Screening Results.** Shown are the results of representative cytotoxicity assays for the 94 Hammy genes screened in this study. Each strain was spotted in triplicate alongside *M. smegmatis*/pExTra-Fruitloop52 (+) and pExTra-Fruitloop52I70S (-) control strains on 7H11 Kan supplemented with 0, 10, or 100 ng/ml aTc. In all experiments, $10^{-1}$ to $10^{-5}$ dilutions are shown; in some experiments the undiluted sample was also spotted. Plates were monitored over 3 or 4 days at 37 ºC, with results shown to best illustrate effects on colony color and size. Colony color was scored using the indicated key.

### Gene *1*; Score *0*

Images taken after 3 days at 37 ºC

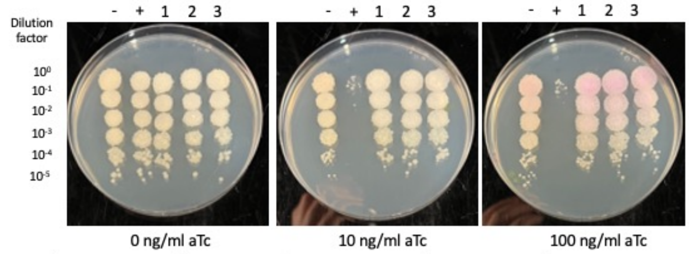


| Lane | Plasmid name | Gene name, replicate | Toxic/Non-toxic | Colony color on 100 ng/ml aTc plate* |
|---|---|---|---|---|
| - Non-toxic control | pExTra03 | Fruitloop *52* mutant | Non-toxic | + |
| + Toxic control | pExTra02 | Fruitloop *52* | Toxic | - |
| 1 | pExTra-Hammy1 | Hammy *1* replicate 1 | Non-toxic | ++ |
| 2 | pExTra-Hammy1 | Hammy *1* replicate 2 | Non-toxic | ++ |
| 3 | pExTra-Hammy1 | Hammy *1* replicate 3 | Non-toxic | ++ |

*Key: NG (no growth) - (no pink color) +(faint pink color) ++(obvious pink color) +++ (dark pink color)

### Gene *2*; Score *0*

Images taken after 4 days at 37 ºC

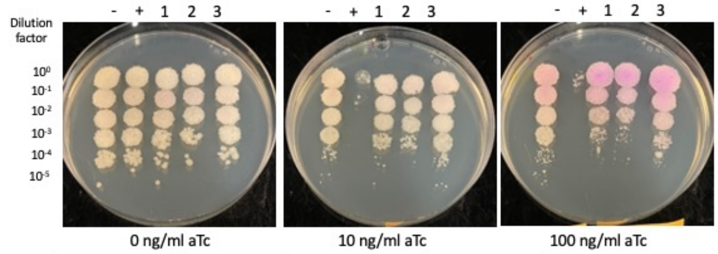


| Lane | Plasmid name | Gene name, replicate | Toxic/Non-toxic | Colony color on 100 ng/ml aTc plate* |
|---|---|---|---|---|
| - Non-toxic control | pExTra03 | Fruitloop *52* mutant | Non-toxic | + |
| + Toxic control | pExTra02 | Fruitloop *52* | Toxic | - |
| 1 | pExTra-Hammy2 | Hammy *2* replicate 1 | Non-toxic | ++ |
| 2 | pExTra-Hammy2 | Hammy *2* replicate 2 | Non-toxic | ++ |
| 3 | pExTra-Hammy2 | Hammy *2* replicate 3 | Non-toxic | ++ |

*Key: NG (no growth) - (no pink color) +(faint pink color) ++(obvious pink color) +++ (dark pink color)

### Gene *3*; Score *0*

Images taken after 4 days at 37 ºC

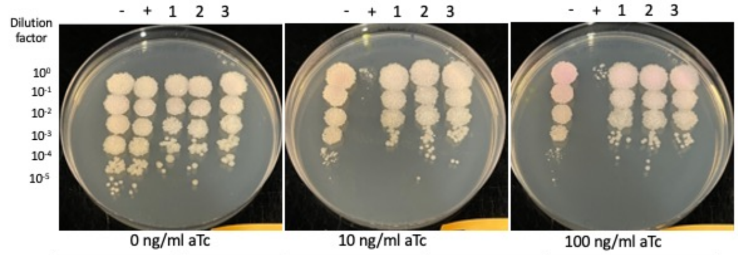


| Lane | Plasmid name | Gene name, replicate | Toxic/Non-toxic | Colony color on 100 ng/ml aTc plate* |
|---|---|---|---|---|
| - Non-toxic control | pExTra03 | Fruitloop *52* mutant | Non-toxic | + |
| + Toxic control | pExTra02 | Fruitloop *52* | Toxic | - |
| 1 | pExTra-Hammy3 | Hammy *3* replicate 1 | Non-toxic | + |
| 2 | pExTra-Hammy3 | Hammy *3* replicate 2 | Non-toxic | + |
| 3 | pExTra-Hammy3 | Hammy *3* replicate 3 | Non-toxic | + |

*Key: NG (no growth) - (no pink color) +(faint pink color) ++(obvious pink color) +++ (dark pink color)

### Gene *5*; Score *0*

Images taken after 4 days at 37 ºC

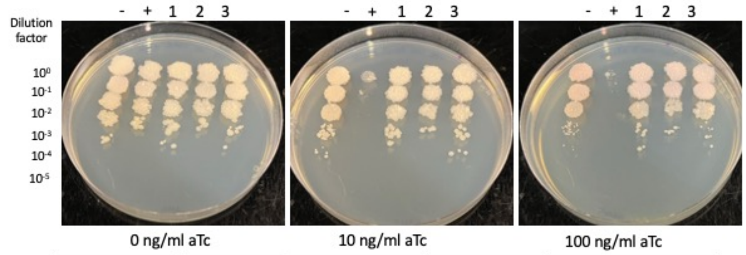


| Lane | Plasmid name | Gene name, replicate | Toxic/Non-toxic | Colony color on 100 ng/ml aTc plate* |
|---|---|---|---|---|
| - Non-toxic control | pExTra03 | Fruitloop *52* mutant | Non-toxic | + |
| + Toxic control | pExTra02 | Fruitloop *52* | Toxic | - |
| 1 | pExTra-Hammy5 | Hammy *5* replicate 1 | Non-toxic | + |
| 2 | pExTra-Hammy5 | Hammy *5* replicate 2 | Non-toxic | + |
| 3 | pExTra-Hammy5 | Hammy *5* replicate 3 | Non-toxic | + |

*Key: NG (no growth) - (no pink color) +(faint pink color) ++(obvious pink color) +++ (dark pink color)

### Gene *6*; Score *0*

Images taken after 4 days at 37 ºC

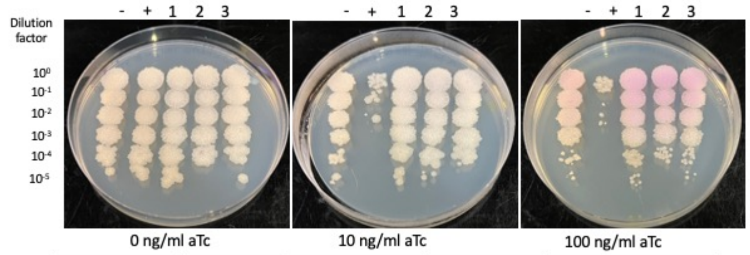


| Lane | Plasmid name | Gene name, replicate | Toxic/Non-toxic | Colony color on 100 ng/ml aTc plate* |
|---|---|---|---|---|
| - Non-toxic control | pExTra03 | Fruitloop *52* mutant | Non-toxic | + |
| + Toxic control | pExTra02 | Fruitloop *52* | Toxic | - |
| 1 | pExTra-Hammy6 | Hammy *6* replicate 1 | Non-toxic | ++ |
| 2 | pExTra-Hammy6 | Hammy *6* replicate 2 | Non-toxic | ++ |
| 3 | pExTra-Hammy6 | Hammy *6* replicate 3 | Non-toxic | ++ |

*Key: NG (no growth) - (no pink color) +(faint pink color) ++(obvious pink color) +++ (dark pink color)

### Gene *7*; Score *0*

Images taken after 3 days at 37 ºC

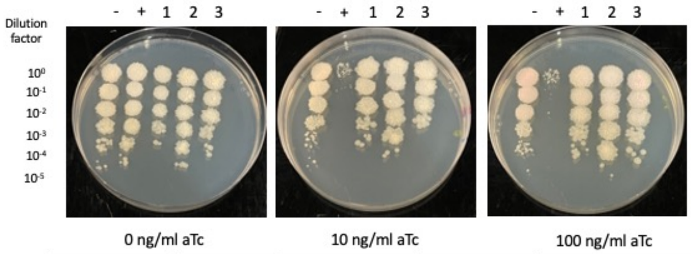


| Lane | Plasmid name | Gene name, replicate | Toxic/Non-toxic | Colony color on 100 ng/ml aTc plate* |
|---|---|---|---|---|
| - Non-toxic control | pExTra03 | Fruitloop *52* mutant | Non-toxic | + |
| + Toxic control | pExTra02 | Fruitloop *52* | Toxic | - |
| 1 | pExTra-Hammy7 | Hammy *7* replicate 1 | Non-toxic | - |
| 2 | pExTra-Hammy7 | Hammy *7* replicate 2 | Non-toxic | - |
| 3 | pExTra-Hammy7 | Hammy *7* replicate 3 | Non-toxic | - |

*Key: NG (no growth) - (no pink color) +(faint pink color) ++(obvious pink color) +++ (dark pink color)

### Gene *8*; Score *0*

Images taken after 3 days at 37 ºC

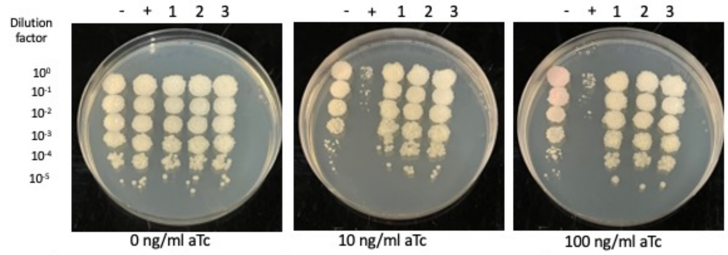


| Lane | Plasmid name | Gene name, replicate | Toxic/Non-toxic | Colony color on 100 ng/ml aTc plate* |
|---|---|---|---|---|
| - Non-toxic control | pExTra03 | Fruitloop *52* mutant | Non-toxic | + |
| + Toxic control | pExTra02 | Fruitloop *52* | Toxic | - |
| 1 | pExTra-Hammy8 | Hammy *8* replicate 1 | Non-toxic | - |
| 2 | pExTra-Hammy8 | Hammy *8* replicate 2 | Non-toxic | - |
| 3 | pExTra-Hammy8 | Hammy *8* replicate 3 | Non-toxic | - |

*Key: NG (no growth) - (no pink color) +(faint pink color) ++(obvious pink color) +++ (dark pink color)

### Gene *9*; Score *3*

Images taken after 3 days at 37 ºC

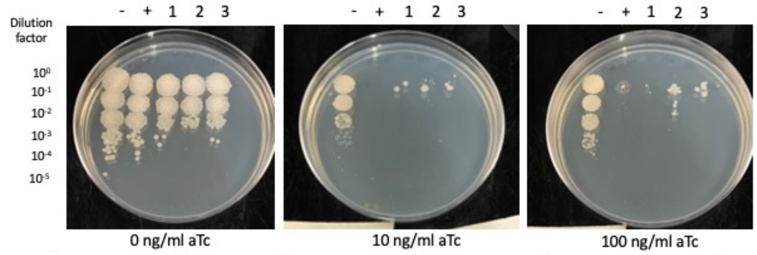


| Lane | Plasmid name | Gene name, replicate | Toxic/Non-toxic | Colony color on 100 ng/ml aTc plate* |
|---|---|---|---|---|
| - Non-toxic control | pExTra03 | Fruitloop *52* mutant | Non-toxic | + |
| + Toxic control | pExTra02 | Fruitloop *52* | Toxic | - |
| 1 | pExTra-Hammy9 | Hammy *9* replicate 1 | Toxic | -/NG |
| 2 | pExTra-Hammy9 | Hammy *9* replicate 2 | Toxic | -/NG |
| 3 | pExTra-Hammy9 | Hammy *9* replicate 3 | Toxic | -/NG |

*Key: NG (no growth) - (no pink color) +(faint pink color) ++(obvious pink color) +++ (dark pink color)

Images taken after 4 days at 37 ºC

### Gene *10*; Score *0*

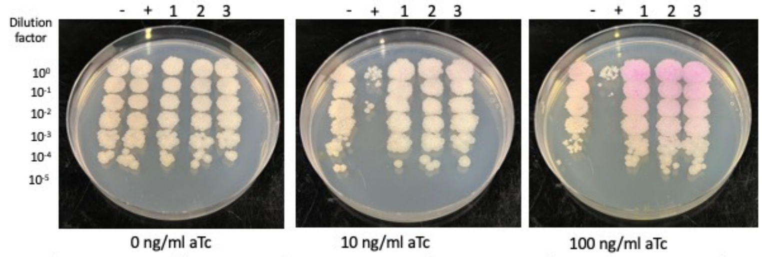


| Lane | Plasmid name | Gene name, replicate | Toxic/Non-toxic | Colony color on 100 ng/ml aTc plate* |
|---|---|---|---|---|
| - Non-toxic control | pExTra03 | Fruitloop *52* mutant | Non-toxic | + |
| + Toxic control | pExTra02 | Fruitloop *52* | Toxic | - |
| 1 | pExTra-Hammy10 | Hammy *10* replicate 1 | Non-toxic | +++ |
| 2 | pExTra-Hammy10 | Hammy *10* replicate 2 | Non-toxic | +++ |
| 3 | pExTra-Hammy10 | Hammy *10* replicate 3 | Non-toxic | +++ |

*Key: NG (no growth) - (no pink color) +(faint pink color) ++(obvious pink color) +++ (dark pink color)

Images taken after 3 days at 37 ºC

### Gene *11*; Score *0*

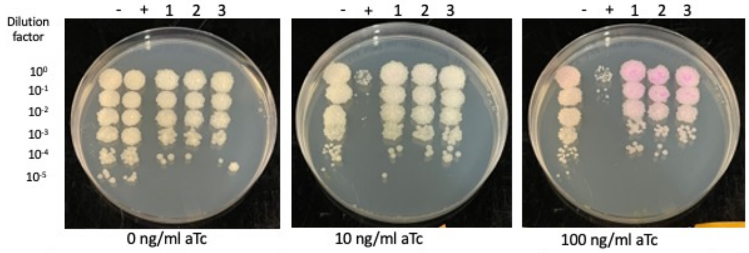


| Lane | Plasmid name | Gene name, replicate | Toxic/Non-toxic | Colony color on 100 ng/ml aTc plate* |
|---|---|---|---|---|
| - Non-toxic control | pExTra03 | Fruitloop *52* mutant | Non-toxic | + |
| + Toxic control | pExTra02 | Fruitloop *52* | Toxic | - |
| 1 | pExTra-Hammy11 | Hammy *11* replicate 1 | Non-toxic | ++ |
| 2 | pExTra-Hammy11 | Hammy *11* replicate 2 | Non-toxic | ++ |
| 3 | pExTra-Hammy11 | Hammy *11* replicate 3 | Non-toxic | ++ |

*Key: NG (no growth) - (no pink color) +(faint pink color) ++(obvious pink color) +++ (dark pink color)

Images taken after 4 days at 37 ºC

### Gene *12*; Score *1*

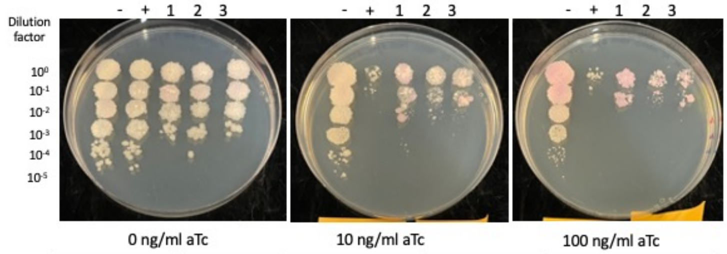


| Lane | Plasmid name | Gene name, replicate | Toxic/Non-toxic | Colony color on 100 ng/ml aTc plate* |
|---|---|---|---|---|
| - Non-toxic control | pExTra03 | Fruitloop *52* mutant | Non-toxic | ++ |
| + Toxic control | pExTra02 | Fruitloop *52* | Toxic | - |
| 1 | pExTra-Hammy12 | Hammy *12* replicate 1 | Toxic | ++ |
| 2 | pExTra-Hammy12 | Hammy *12* replicate 2 | Toxic | ++ |
| 3 | pExTra-Hammy12 | Hammy *12* replicate 3 | Toxic | ++ |

*Key: NG (no growth) - (no pink color) +(faint pink color) ++(obvious pink color) +++ (dark pink color)

Images taken after 4 days at 37 ºC

### Gene *13*; Score *0*

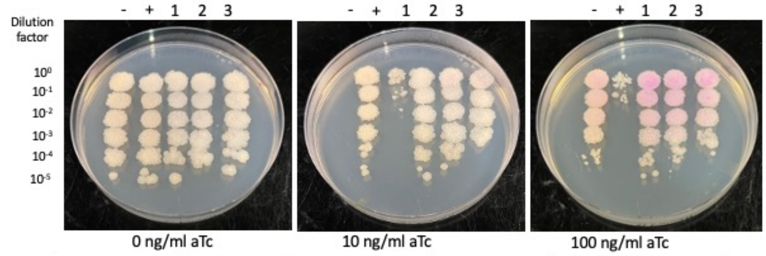


| Lane | Plasmid name | Gene name, replicate | Toxic/Non-toxic | Colony color on 100 ng/ml aTc plate* |
|---|---|---|---|---|
| - Non-toxic control | pExTra03 | Fruitloop *52* mutant | Non-toxic | + |
| + Toxic control | pExTra02 | Fruitloop *52* | Toxic | - |
| 1 | pExTra-Hammy13 | Hammy *13* replicate 1 | Non-toxic | ++ |
| 2 | pExTra-Hammy13 | Hammy *13* replicate 2 | Non-toxic | ++ |
| 3 | pExTra-Hammy13 | Hammy *13* replicate 3 | Non-toxic | ++ |

*Key: NG (no growth) - (no pink color) +(faint pink color) ++(obvious pink color) +++ (dark pink color)

Images taken after 4 days at 37 ºC

### Gene *14*; Score *0*

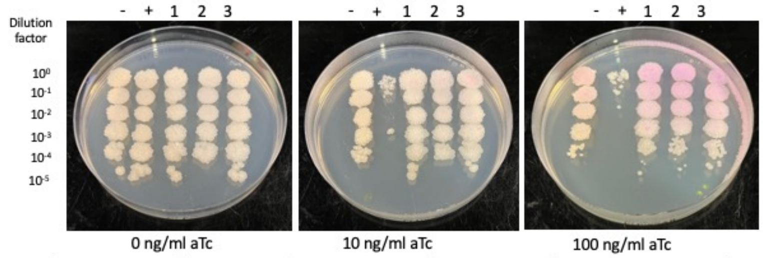


| Lane | Plasmid name | Gene name, replicate | Toxic/Non-toxic | Colony color on 100 ng/ml aTc plate* |
|---|---|---|---|---|
| - Non-toxic control | pExTra03 | Fruitloop *52* mutant | Non-toxic | + |
| + Toxic control | pExTra02 | Fruitloop *52* | Toxic | - |
| 1 | pExTra-Hammy14 | Hammy *14* replicate 1 | Non-toxic | ++ |
| 2 | pExTra-Hammy14 | Hammy *14* replicate 2 | Non-toxic | ++ |
| 3 | pExTra-Hammy14 | Hammy *14* replicate 3 | Non-toxic | ++ |

*Key: NG (no growth) - (no pink color) +(faint pink color) ++(obvious pink color) +++ (dark pink color)

Images taken after 4 days at 37 ºC

### Gene *15*; Score *0*

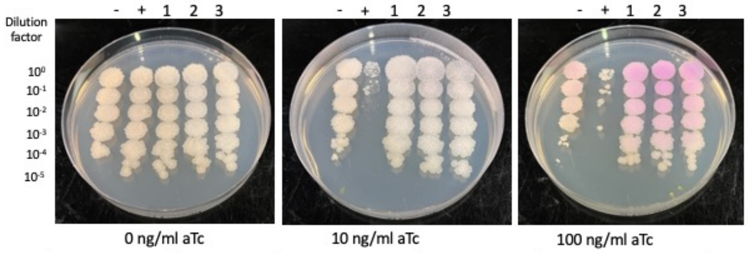


| Lane | Plasmid name | Gene name, replicate | Toxic/Non-toxic | Colony color on 100 ng/ml aTc plate* |
|---|---|---|---|---|
| - Non-toxic control | pExTra03 | Fruitloop *52* mutant | Non-toxic | + |
| + Toxic control | pExTra02 | Fruitloop *52* | Toxic | - |
| 1 | pExTra-Hammy15 | Hammy *15* replicate 1 | Non-toxic | +++ |
| 2 | pExTra-Hammy15 | Hammy *15* replicate 2 | Non-toxic | +++ |
| 3 | pExTra-Hammy15 | Hammy *15* replicate 3 | Non-toxic | +++ |

*Key: NG (no growth) - (no pink color) +(faint pink color) ++(obvious pink color) +++ (dark pink color)

Images taken after 4 days at 37 ºC

### Gene *16*; Score *0*

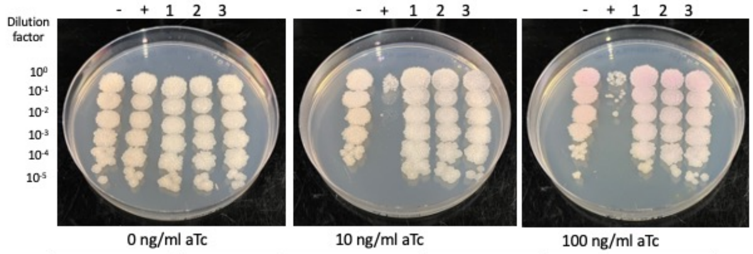


| Lane | Plasmid name | Gene name, replicate | Toxic/Non-toxic | Colony color on 100 ng/ml aTc plate* |
|---|---|---|---|---|
| - Non-toxic control | pExTra03 | Fruitloop *52* mutant | Non-toxic | + |
| + Toxic control | pExTra02 | Fruitloop *52* | Toxic | - |
| 1 | pExTra-Hammy16 | Hammy *16* replicate 1 | Non-toxic | + |
| 2 | pExTra-Hammy16 | Hammy *16* replicate 2 | Non-toxic | + |
| 3 | pExTra-Hammy16 | Hammy *16* replicate 3 | Non-toxic | + |

*Key: NG (no growth) - (no pink color) +(faint pink color) ++(obvious pink color) +++ (dark pink color)

Images taken after 4 days at 37 ºC

### Gene *17*; Score *0*

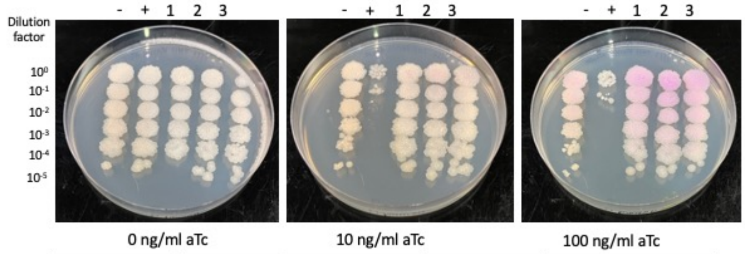


| Lane | Plasmid name | Gene name, replicate | Toxic/Non-toxic | Colony color on 100 ng/ml aTc plate* |
|---|---|---|---|---|
| - Non-toxic control | pExTra03 | Fruitloop *52* mutant | Non-toxic | + |
| + Toxic control | pExTra02 | Fruitloop *52* | Toxic | - |
| 1 | pExTra-Hammy17 | Hammy *17* replicate 1 | Non-toxic | ++ |
| 2 | pExTra-Hammy17 | Hammy *17* replicate 2 | Non-toxic | ++ |
| 3 | pExTra-Hammy17 | Hammy *17* replicate 3 | Non-toxic | ++ |

*Key: NG (no growth) - (no pink color) +(faint pink color) ++(obvious pink color) +++ (dark pink color)

Images taken after 4 days at 37 ºC

Gene *18*; Score *0*

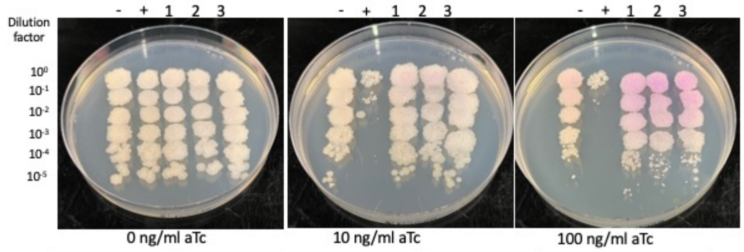


| Lane | Plasmid name | Gene name, replicate | Toxic/Non-toxic | Colony color on 100 ng/ml aTc plate* |
|---|---|---|---|---|
| - Non-toxic control | pExTra03 | Fruitloop *52* mutant | Non-toxic | + |
| + Toxic control | pExTra02 | Fruitloop *52* | Toxic | - |
| 1 | pExTra-Hammy18 | Hammy *18* replicate 1 | Non-toxic | ++ |
| 2 | pExTra-Hammy18 | Hammy *18* replicate 2 | Non-toxic | ++ |
| 3 | pExTra-Hammy18 | Hammy *18* replicate 3 | Non-toxic | ++ |

*Key: NG (no growth) - (no pink color) +(faint pink color) ++(obvious pink color) +++ (dark pink color)

Images taken after 3 days at 37 ºC

Gene *20*; Score 3

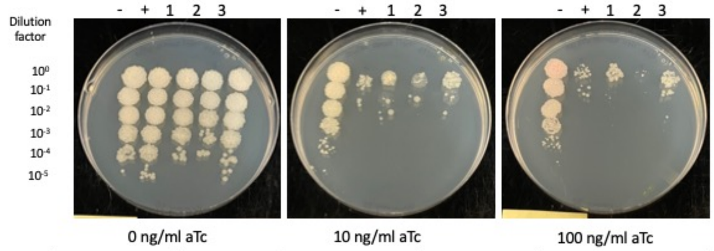


| Lane | Plasmid name | Gene name, replicate | Toxic/Non-toxic | Colony color on 100 ng/ml aTc plate* |
|---|---|---|---|---|
| - Non-toxic control | pExTra03 | Fruitloop *52* mutant | Non-toxic | + |
| + Toxic control | pExTra02 | Fruitloop *52* | Toxic | - |
| 1 | pExTra-Hammy20 | Hammy *20* replicate 1 | Toxic | - |
| 2 | pExTra-Hammy20 | Hammy *20* replicate 2 | Toxic | -/NG |
| 3 | pExTra-Hammy20 | Hammy *20* replicate 3 | Toxic | - |

*Key: NG (no growth) - (no pink color) +(faint pink color) ++(obvious pink color) +++ (dark pink color)

Images taken after 4 days at 37 ºC

Gene *21*; Score *0*

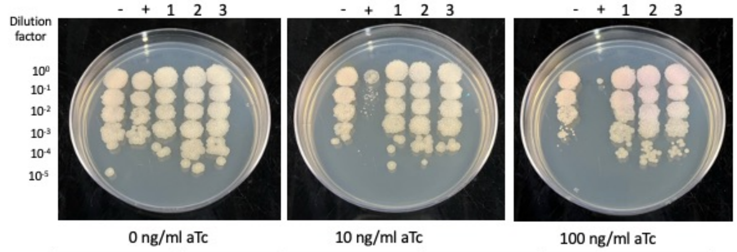


| Lane | Plasmid name | Gene name, replicate | Toxic/Non-toxic | Colony color on 100 ng/ml aTc plate* |
|---|---|---|---|---|
| - Non-toxic control | pExTra03 | Fruitloop *52* mutant | Non-toxic | + |
| + Toxic control | pExTra02 | Fruitloop *52* | Toxic | - |
| 1 | pExTra-Hammy21 | Hammy *21* replicate 1 | Non-toxic | + |
| 2 | pExTra-Hammy21 | Hammy *21* replicate 2 | Non-toxic | + |
| 3 | pExTra-Hammy21 | Hammy *21* replicate 3 | Non-toxic | + |

*Key: NG (no growth) - (no pink color) +(faint pink color) ++(obvious pink color) +++ (dark pink color)

Images taken after 4 days at 37 ºC

Gene *22*; Score *0*

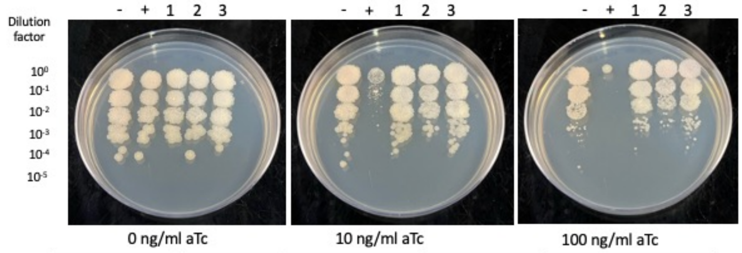


| Lane | Plasmid name | Gene name, replicate | Toxic/Non-toxic | Colony color on 100 ng/ml aTc plate* |
|---|---|---|---|---|
| - Non-toxic control | pExTra03 | Fruitloop *52* mutant | Non-toxic | - |
| + Toxic control | pExTra02 | Fruitloop *52* | Toxic | - |
| 1 | pExTra-Hammy22 | Hammy *22* replicate 1 | Non-toxic | - |
| 2 | pExTra-Hammy22 | Hammy *22* replicate 2 | Non-toxic | - |
| 3 | pExTra-Hammy22 | Hammy *22* replicate 3 | Non-toxic | - |

*Key: NG (no growth) - (no pink color) +(faint pink color) ++(obvious pink color) +++ (dark pink color)

Images taken after 4 days at 37 ºC

Gene *23*; Score *0*

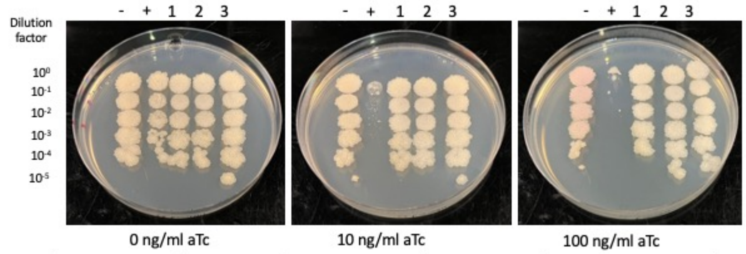


| Lane | Plasmid name | Gene name, replicate | Toxic/Non-toxic | Colony color on 100 ng/ml aTc plate* |
|---|---|---|---|---|
| - Non-toxic control | pExTra03 | Fruitloop *52* mutant | Non-toxic | + |
| + Toxic control | pExTra02 | Fruitloop *52* | Toxic | - |
| 1 | pExTra-Hammy23 | Hammy *23* replicate 1 | Non-toxic | - |
| 2 | pExTra-Hammy23 | Hammy *23* replicate 2 | Non-toxic | - |
| 3 | pExTra-Hammy23 | Hammy *23* replicate 3 | Non-toxic | - |

*Key: NG (no growth) - (no pink color) +(faint pink color) ++(obvious pink color) +++ (dark pink color)

Images taken after 4 days at 37 ºC

Gene *24*; Score *0*

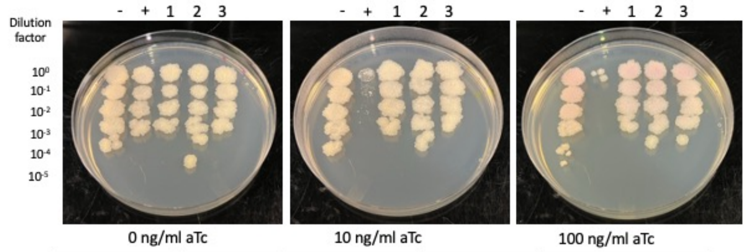


| Lane | Plasmid name | Gene name, replicate | Toxic/Non-toxic | Colony color on 100 ng/ml aTc plate* |
|---|---|---|---|---|
| - Non-toxic control | pExTra03 | Fruitloop *52* mutant | Non-toxic | + |
| + Toxic control | pExTra02 | Fruitloop *52* | Toxic | - |
| 1 | pExTra-Hammy24 | Hammy *24* replicate 1 | Non-toxic | + |
| 2 | pExTra-Hammy24 | Hammy *24* replicate 2 | Non-toxic | + |
| 3 | pExTra-Hammy24 | Hammy *24* replicate 3 | Non-toxic | + |

*Key: NG (no growth) - (no pink color) +(faint pink color) ++(obvious pink color) +++ (dark pink color)

Images taken after 4 days at 37 ºC

Gene *25*; Score *0*

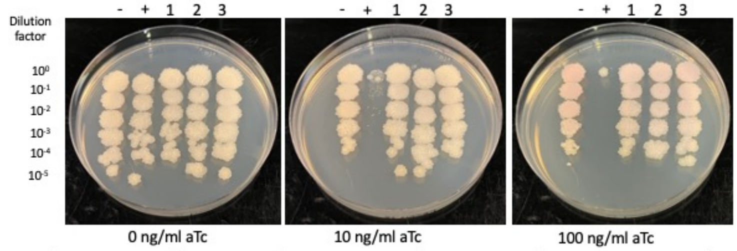


| Lane | Plasmid name | Gene name, replicate | Toxic/Non-toxic | Colony color on 100 ng/ml aTc plate* |
|---|---|---|---|---|
| - Non-toxic control | pExTra03 | Fruitloop *52* mutant | Non-toxic | + |
| + Toxic control | pExTra02 | Fruitloop *52* | Toxic | - |
| 1 | pExTra-Hammy25 | Hammy *25* replicate 1 | Non-toxic | + |
| 2 | pExTra-Hammy25 | Hammy *25* replicate 2 | Non-toxic | + |
| 3 | pExTra-Hammy25 | Hammy *25* replicate 3 | Non-toxic | + |

*Key: NG (no growth) - (no pink color) +(faint pink color) ++(obvious pink color) +++ (dark pink color)

Images taken after 3 days at 37 ºC

Gene *26*; Score *0*

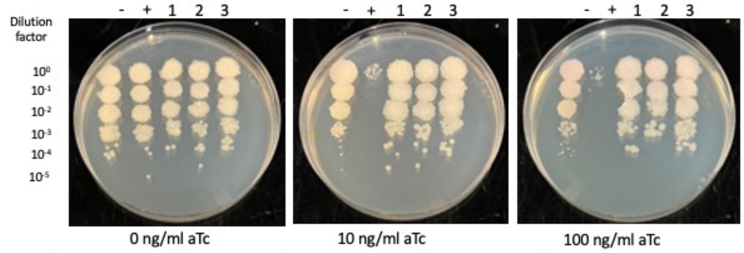


| Lane | Plasmid name | Gene name, replicate | Toxic/Non-toxic | Colony color on 100 ng/ml aTc plate* |
|---|---|---|---|---|
| - Non-toxic control | pExTra03 | Fruitloop *52* mutant | Non-toxic | + |
| + Toxic control | pExTra02 | Fruitloop *52* | Toxic | - |
| 1 | pExTra-Hammy26 | Hammy *26* replicate 1 | Non-toxic | + |
| 2 | pExTra-Hammy26 | Hammy *26* replicate 2 | Non-toxic | + |
| 3 | pExTra-Hammy26 | Hammy *26* replicate 3 | Non-toxic | + |

*Key: NG (no growth) - (no pink color) +(faint pink color) ++(obvious pink color) +++ (dark pink color)

### Gene *27*; Score *0*

Images taken after 3 days at 37 ºC

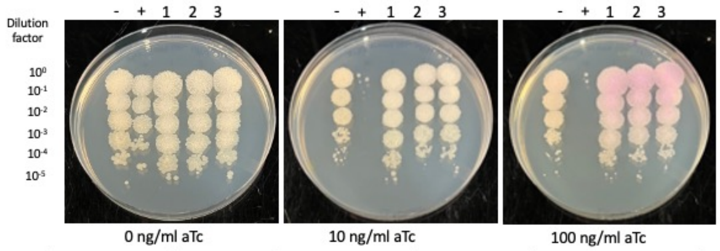


| Lane | Plasmid name | Gene name, replicate | Toxic/Non-toxic | Colony color on 100 ng/ml aTc plate* |
|---|---|---|---|---|
| - Non-toxic control | pExTra03 | Fruitloop *52* mutant | Non-toxic | + |
| + Toxic control | pExTra02 | Fruitloop *52* | Toxic | -/NG |
| 1 | pExTra-Hammy27 | Hammy *27* replicate 1 | Non-toxic | ++ |
| 2 | pExTra-Hammy27 | Hammy *27* replicate 2 | Non-toxic | ++ |
| 3 | pExTra-Hammy27 | Hammy *27* replicate 3 | Non-toxic | ++ |

*Key: NG (no growth) - (no pink color) +(faint pink color) ++(obvious pink color) +++ (dark pink color)

### Gene *28*; Score *0*

Images taken after 4 days at 37 ºC

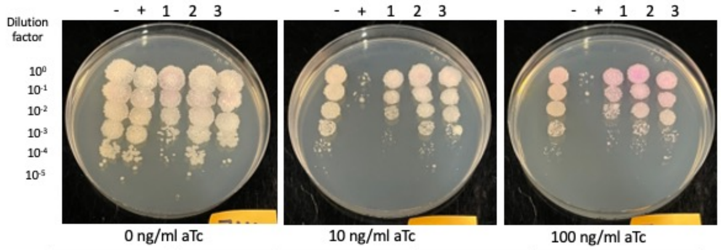


| Lane | Plasmid name | Gene name, replicate | Toxic/Non-toxic | Colony color on 100 ng/ml aTc plate* |
|---|---|---|---|---|
| - Non-toxic control | pExTra03 | Fruitloop *52* mutant | Non-toxic | + |
| + Toxic control | pExTra02 | Fruitloop *52* | Toxic | -/NG |
| 1 | pExTra-Hammy28 | Hammy *28* replicate 1 | Non-toxic | ++ |
| 2 | pExTra-Hammy28 | Hammy *28* replicate 2 | Non-toxic | ++ |
| 3 | pExTra-Hammy28 | Hammy *28* replicate 3 | Non-toxic | ++ |

*Key: NG (no growth) - (no pink color) +(faint pink color) ++(obvious pink color) +++ (dark pink color)

### Gene *29*; Score *1*

Images taken after 3 days at 37 ºC

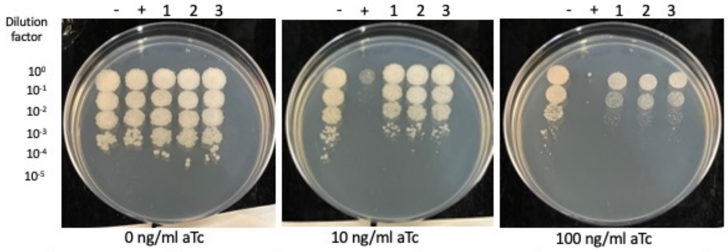


| Lane | Plasmid name | Gene name, replicate | Toxic/Non-toxic | Colony color on 100 ng/ml aTc plate* |
|---|---|---|---|---|
| - Non-toxic control | pExTra03 | Fruitloop *52* mutant | Non-toxic | + |
| + Toxic control | pExTra02 | Fruitloop *52* | Toxic | -/NG |
| 1 | pExTra-Hammy29 | Hammy *29* replicate 1 | Toxic | - |
| 2 | pExTra-Hammy29 | Hammy *29* replicate 2 | Toxic | - |
| 3 | pExTra-Hammy29 | Hammy *29* replicate 3 | Toxic | - |

*Key: NG (no growth) - (no pink color) +(faint pink color) ++(obvious pink color) +++ (dark pink color)

### Gene *30*; Score *0*

Images taken after 4 days at 37 ºC

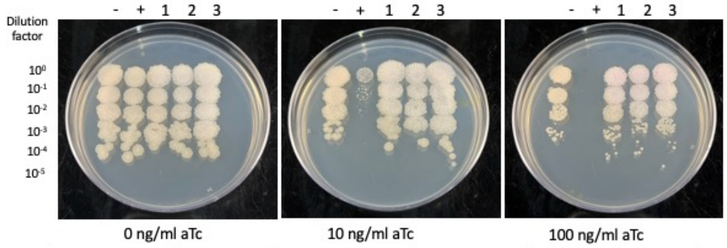


| Lane | Plasmid name | Gene name, replicate | Toxic/Non-toxic | Colony color on 100 ng/ml aTc plate* |
|---|---|---|---|---|
| - Non-toxic control | pExTra03 | Fruitloop *52* mutant | Non-toxic | - |
| + Toxic control | pExTra02 | Fruitloop *52* | Toxic | NG |
| 1 | pExTra-Hammy30 | Hammy *30* replicate 1 | Non-toxic | + |
| 2 | pExTra-Hammy30 | Hammy *30* replicate 2 | Non-toxic | + |
| 3 | pExTra-Hammy30 | Hammy *30* replicate 3 | Non-toxic | + |

*Key: NG (no growth) - (no pink color) +(faint pink color) ++(obvious pink color) +++ (dark pink color)

### Gene *31*; Score *0*

Images taken after 4 days at 37 ºC

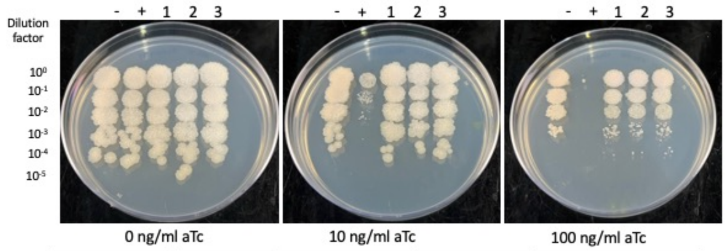


| Lane | Plasmid name | Gene name, replicate | Toxic/Non-toxic | Colony color on 100 ng/ml aTc plate* |
|---|---|---|---|---|
| - Non-toxic control | pExTra03 | Fruitloop *52* mutant | Non-toxic | - |
| + Toxic control | pExTra02 | Fruitloop *52* | Toxic | -/NG |
| 1 | pExTra-Hammy31 | Hammy *31* replicate 1 | Non-toxic | - |
| 2 | pExTra-Hammy31 | Hammy *31* replicate 2 | Non-toxic | - |
| 3 | pExTra-Hammy31 | Hammy *31* replicate 3 | Non-toxic | - |

*Key: NG (no growth) - (no pink color) +(faint pink color) ++(obvious pink color) +++ (dark pink color)

### Gene *32*; Score *2*

Images taken after 4 days at 37 ºC

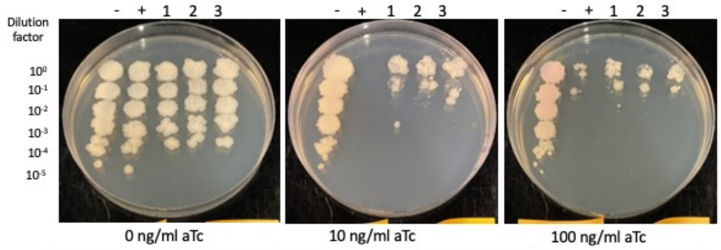


| Lane | Plasmid name | Gene name, replicate | Toxic/Non-toxic | Colony color on 100 ng/ml aTc plate* |
|---|---|---|---|---|
| - Non-toxic control | pExTra03 | Fruitloop *52* mutant | Non-toxic | + |
| + Toxic control | pExTra02 | Fruitloop *52* | Toxic | - |
| 1 | pExTra-Hammy32 | Hammy *32* replicate 1 | Toxic | - |
| 2 | pExTra-Hammy32 | Hammy *32* replicate 2 | Toxic | - |
| 3 | pExTra-Hammy32 | Hammy *32* replicate 3 | Toxic | - |

*Key: NG (no growth) - (no pink color) +(faint pink color) ++(obvious pink color) +++ (dark pink color)

### Gene *33*; Score *0*

Images taken after 4 days at 37 ºC

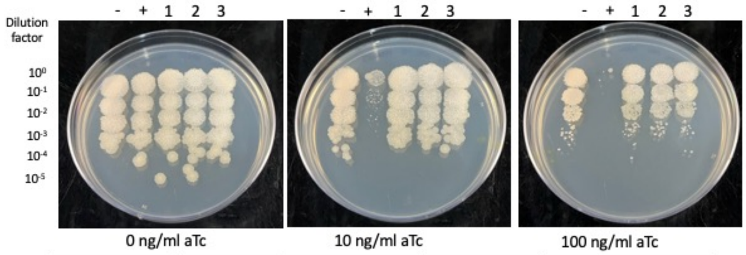


| Lane | Plasmid name | Gene name, replicate | Toxic/Non-toxic | Colony color on 100 ng/ml aTc plate* |
|---|---|---|---|---|
| - Non-toxic control | pExTra03 | Fruitloop *52* mutant | Non-toxic | - |
| + Toxic control | pExTra02 | Fruitloop *52* | Toxic | - |
| 1 | pExTra-Hammy33 | Hammy *33* replicate 1 | Non-toxic | - |
| 2 | pExTra-Hammy33 | Hammy *33* replicate 2 | Non-toxic | - |
| 3 | pExTra-Hammy33 | Hammy *33* replicate 3 | Non-toxic | - |

*Key: NG (no growth) - (no pink color) +(faint pink color) ++(obvious pink color) +++ (dark pink color)

### Gene *34*; Score *3*

Images taken after 3 days at 37 ºC

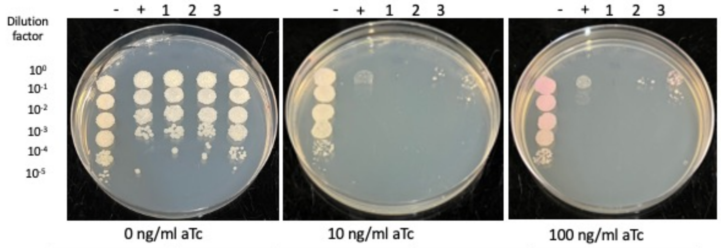


| Lane | Plasmid name | Gene name, replicate | Toxic/Non-toxic | Colony color on 100 ng/ml aTc plate* |
|---|---|---|---|---|
| - Non-toxic control | pExTra03 | Fruitloop *52* mutant | Non-toxic | ++ |
| + Toxic control | pExTra02 | Fruitloop *52* | Toxic | - |
| 1 | pExTra-Hammy34 | Hammy *34* replicate 1 | Toxic | -/NG |
| 2 | pExTra-Hammy34 | Hammy *34* replicate 2 | Toxic | -/NG |
| 3 | pExTra-Hammy34 | Hammy *34* replicate 3 | Toxic | + |

*Key: NG (no growth) - (no pink color) +(faint pink color) ++(obvious pink color) +++ (dark pink color)

Images taken after 3 days at 37 ºC

### Gene 35; Score 0

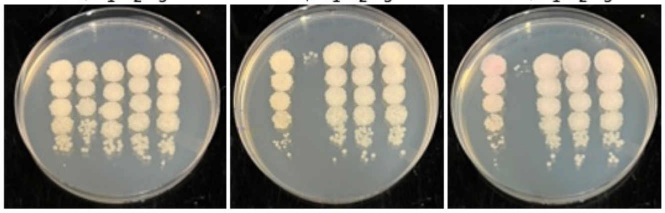


| Lane | Plasmid name | Gene name, replicate | Toxic/Non-toxic | Colony color on 100 ng/ml aTc plate* |
|---|---|---|---|---|
| - Non-toxic control | pExTra03 | Fruitloop *52* mutant | Non-toxic | + |
| + Toxic control | pExTra02 | Fruitloop *52* | Toxic | -/NG |
| 1 | pExTra-Hammy35 | Hammy *35* replicate 1 | Non-toxic | - |
| 2 | pExTra-Hammy35 | Hammy *35* replicate 2 | Non-toxic | - |
| 3 | pExTra-Hammy35 | Hammy *35* replicate 3 | Non-toxic | - |

*Key: NG (no growth) - (no pink color) +(faint pink color) ++(obvious pink color) +++ (dark pink color)

Images taken after 3 days at 37 ºC

### Gene 36; Score 1

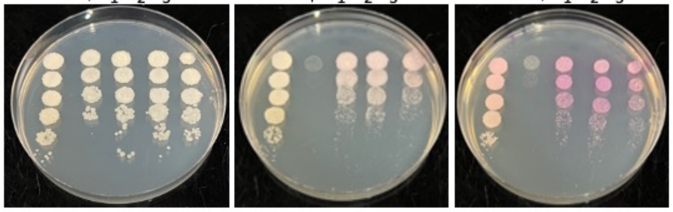


| Lane | Plasmid name | Gene name, replicate | Toxic/Non-toxic | Colony color on 100 ng/ml aTc plate* |
|---|---|---|---|---|
| - Non-toxic control | pExTra03 | Fruitloop *52* mutant | Non-toxic | ++ |
| + Toxic control | pExTra02 | Fruitloop *52* | Toxic | - |
| 1 | pExTra-Hammy36 | Hammy *36* replicate 1 | Toxic | +++ |
| 2 | pExTra-Hammy36 | Hammy *36* replicate 2 | Toxic | +++ |
| 3 | pExTra-Hammy36 | Hammy *36* replicate 3 | Toxic | +++ |

*Key: NG (no growth) - (no pink color) +(faint pink color) ++(obvious pink color) +++ (dark pink color)

Images taken after 4 days at 37 ºC

### Gene 37; Score 0

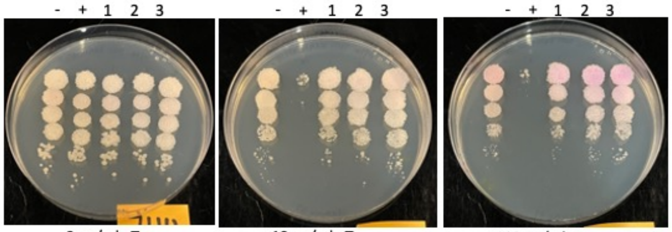


| Lane | Plasmid name | Gene name, replicate | Toxic/Non-toxic | Colony color on 100 ng/ml aTc plate* |
|---|---|---|---|---|
| - Non-toxic control | pExTra03 | Fruitloop *52* mutant | Non-toxic | + |
| + Toxic control | pExTra02 | Fruitloop *52* | Toxic | -/NG |
| 1 | pExTra-Hammy37 | Hammy *37* replicate 1 | Non-toxic | ++ |
| 2 | pExTra-Hammy37 | Hammy *37* replicate 2 | Non-toxic | ++ |
| 3 | pExTra-Hammy37 | Hammy *37* replicate 3 | Non-toxic | ++ |

*Key: NG (no growth) - (no pink color) +(faint pink color) ++(obvious pink color) +++ (dark pink color)

Images taken after 4 days at 37 ºC

### Gene 38; Score 0

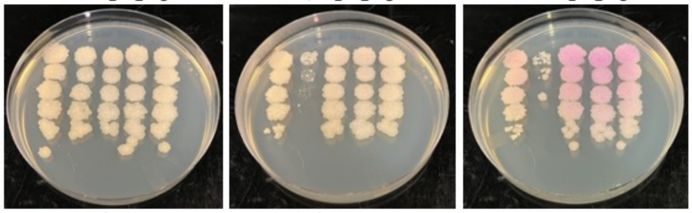


| Lane | Plasmid name | Gene name, replicate | Toxic/Non-toxic | Colony color on 100 ng/ml aTc plate* |
|---|---|---|---|---|
| - Non-toxic control | pExTra03 | Fruitloop *52* mutant | Non-toxic | + |
| + Toxic control | pExTra02 | Fruitloop *52* | Toxic | - |
| 1 | pExTra-Hammy38 | Hammy *38* replicate 1 | Non-toxic | +++ |
| 2 | pExTra-Hammy38 | Hammy *38* replicate 2 | Non-toxic | +++ |
| 3 | pExTra-Hammy38 | Hammy *38* replicate 3 | Non-toxic | +++ |

*Key: NG (no growth) - (no pink color) +(faint pink color) ++(obvious pink color) +++ (dark pink color)

Images taken after 4 days at 37 ºC

### Gene 39; Score 0

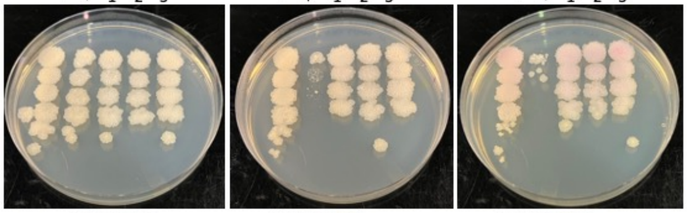


| Lane | Plasmid name | Gene name, replicate | Toxic/Non-toxic | Colony color on 100 ng/ml aTc plate* |
|---|---|---|---|---|
| - Non-toxic control | pExTra03 | Fruitloop *52* mutant | Non-toxic | + |
| + Toxic control | pExTra02 | Fruitloop *52* | Toxic | - |
| 1 | pExTra-Hammy39 | Hammy *39* replicate 1 | Non-toxic | + |
| 2 | pExTra-Hammy39 | Hammy *39* replicate 2 | Non-toxic | + |
| 3 | pExTra-Hammy39 | Hammy *39* replicate 3 | Non-toxic | + |

*Key: NG (no growth) - (no pink color) +(faint pink color) ++(obvious pink color) +++ (dark pink color)

Images taken after 4 days at 37 ºC

### Gene 40; Score 0

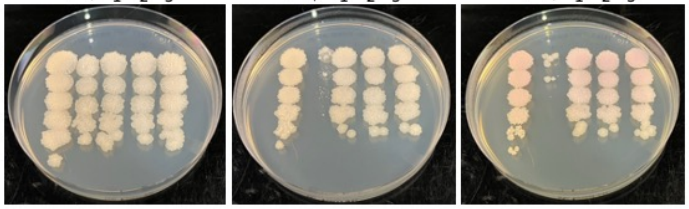


| Lane | Plasmid name | Gene name, replicate | Toxic/Non-toxic | Colony color on 100 ng/ml aTc plate* |
|---|---|---|---|---|
| - Non-toxic control | pExTra03 | Fruitloop *52* mutant | Non-toxic | + |
| + Toxic control | pExTra02 | Fruitloop *52* | Toxic | - |
| 1 | pExTra-Hammy40 | Hammy *40* replicate 1 | Non-toxic | + |
| 2 | pExTra-Hammy40 | Hammy *40* replicate 2 | Non-toxic | + |
| 3 | pExTra-Hammy40 | Hammy *40* replicate 3 | Non-toxic | + |

*Key: NG (no growth) - (no pink color) +(faint pink color) ++(obvious pink color) +++ (dark pink color)

Images taken after 4 days at 37 ºC

### Gene 41; Score 0

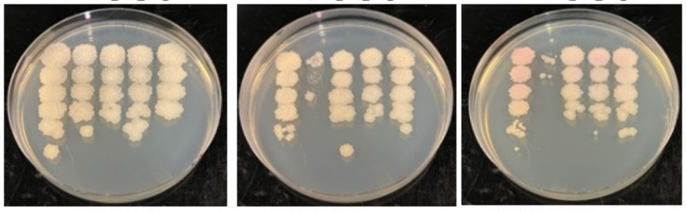


| Lane | Plasmid name | Gene name, replicate | Toxic/Non-toxic | Colony color on 100 ng/ml aTc plate* |
|---|---|---|---|---|
| - Non-toxic control | pExTra03 | Fruitloop *52* mutant | Non-toxic | + |
| + Toxic control | pExTra02 | Fruitloop *52* | Toxic | - |
| 1 | pExTra-Hammy41 | Hammy *41* replicate 1 | Non-toxic | + |
| 2 | pExTra-Hammy41 | Hammy *41* replicate 2 | Non-toxic | + |
| 3 | pExTra-Hammy41 | Hammy *41* replicate 3 | Non-toxic | + |

*Key: NG (no growth) - (no pink color) +(faint pink color) ++(obvious pink color) +++ (dark pink color)

Images taken after 4 days at 37 ºC

### Gene 42; Score 0

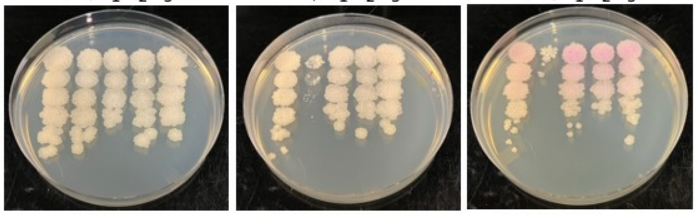


| Lane | Plasmid name | Gene name, replicate | Toxic/Non-toxic | Colony color on 100 ng/ml aTc plate* |
|---|---|---|---|---|
| - Non-toxic control | pExTra03 | Fruitloop *52* mutant | Non-toxic | + |
| + Toxic control | pExTra02 | Fruitloop *52* | Toxic | - |
| 1 | pExTra-Hammy42 | Hammy *42* replicate 1 | Non-toxic | ++ |
| 2 | pExTra-Hammy42 | Hammy *42* replicate 2 | Non-toxic | ++ |
| 3 | pExTra-Hammy42 | Hammy *42* replicate 3 | Non-toxic | ++ |

*Key: NG (no growth) - (no pink color) +(faint pink color) ++(obvious pink color) +++ (dark pink color)

### Gene *43*; Score *0*

Images taken after 4 days at 37 ºC

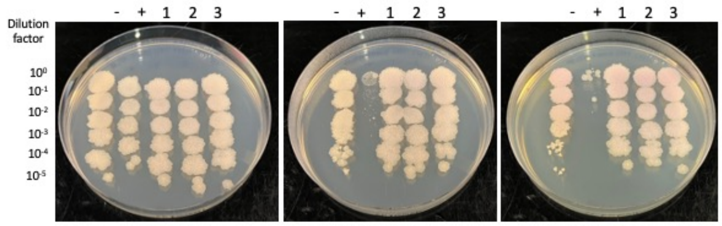


0 ng/ml aTc | 10 ng/ml aTc | 100 ng/ml aTc

| Lane | Plasmid name | Gene name, replicate | Toxic/Non-toxic | Colony color on 100 ng/ml aTc plate* |
|---|---|---|---|---|
| - Non-toxic control | pExTra03 | Fruitloop *52* mutant | Non-toxic | + |
| + Toxic control | pExTra02 | Fruitloop *52* | Toxic | - |
| 1 | pExTra-Hammy43 | Hammy *43* replicate 1 | Non-toxic | + |
| 2 | pExTra-Hammy43 | Hammy *43* replicate 2 | Non-toxic | + |
| 3 | pExTra-Hammy43 | Hammy *43* replicate 3 | Non-toxic | + |

*Key: NG (no growth) - (no pink color) +(faint pink color) ++(obvious pink color) +++ (dark pink color)

### Gene *44*; Score *0*

Images taken after 4 days at 37 ºC

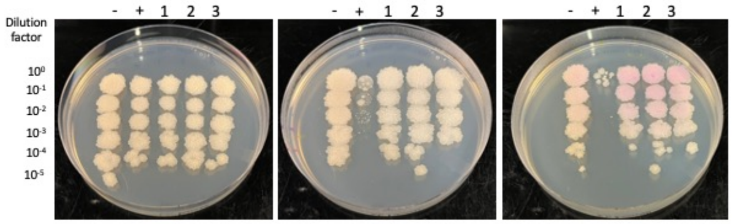


0 ng/ml aTc | 10 ng/ml aTc | 100 ng/ml aTc

| Lane | Plasmid name | Gene name, replicate | Toxic/Non-toxic | Colony color on 100 ng/ml aTc plate* |
|---|---|---|---|---|
| - Non-toxic control | pExTra03 | Fruitloop *52* mutant | Non-toxic | + |
| + Toxic control | pExTra02 | Fruitloop *52* | Toxic | - |
| 1 | pExTra-Hammy44 | Hammy *44* replicate 1 | Non-toxic | ++ |
| 2 | pExTra-Hammy44 | Hammy *44* replicate 2 | Non-toxic | ++ |
| 3 | pExTra-Hammy44 | Hammy *44* replicate 3 | Non-toxic | ++ |

*Key: NG (no growth) - (no pink color) +(faint pink color) ++(obvious pink color) +++ (dark pink color)

### Gene *45*; Score *0*

Images taken after 4 days at 37 ºC

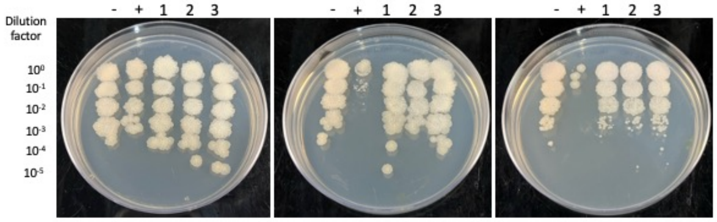


0 ng/ml aTc | 10 ng/ml aTc | 100 ng/ml aTc

| Lane | Plasmid name | Gene name, replicate | Toxic/Non-toxic | Colony color on 100 ng/ml aTc plate* |
|---|---|---|---|---|
| - Non-toxic control | pExTra03 | Fruitloop *52* mutant | Non-toxic | + |
| + Toxic control | pExTra02 | Fruitloop *52* | Toxic | - |
| 1 | pExTra-Hammy45 | Hammy *45* replicate 1 | Non-toxic | - |
| 2 | pExTra-Hammy45 | Hammy *45* replicate 2 | Non-toxic | - |
| 3 | pExTra-Hammy45 | Hammy *45* replicate 3 | Non-toxic | - |

*Key: NG (no growth) - (no pink color) +(faint pink color) ++(obvious pink color) +++ (dark pink color)

### Gene *46*; Score *0*

Images taken after 4 days at 37 ºC

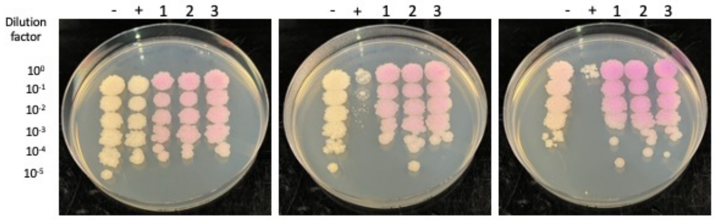


0 ng/ml aTc | 10 ng/ml aTc | 100 ng/ml aTc

| Lane | Plasmid name | Gene name, replicate | Toxic/Non-toxic | Colony color on 100 ng/ml aTc plate* |
|---|---|---|---|---|
| - Non-toxic control | pExTra03 | Fruitloop *52* mutant | Non-toxic | + |
| + Toxic control | pExTra02 | Fruitloop *52* | Toxic | - |
| 1 | pExTra-Hammy46 | Hammy *46* replicate 1 | Non-toxic | +++ |
| 2 | pExTra-Hammy46 | Hammy *46* replicate 2 | Non-toxic | +++ |
| 3 | pExTra-Hammy46 | Hammy *46* replicate 3 | Non-toxic | +++ |

*Key: NG (no growth) - (no pink color) +(faint pink color) ++(obvious pink color) +++ (dark pink color)

### Gene *47*; Score *0*

Images taken after 4 days at 37 ºC

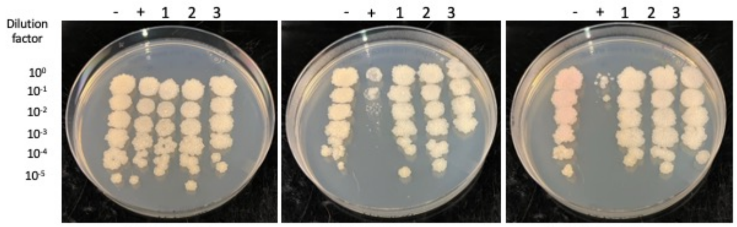


0 ng/ml aTc | 10 ng/ml aTc | 100 ng/ml aTc

| Lane | Plasmid name | Gene name, replicate | Toxic/Non-toxic | Colony color on 100 ng/ml aTc plate* |
|---|---|---|---|---|
| - Non-toxic control | pExTra03 | Fruitloop *52* mutant | Non-toxic | + |
| + Toxic control | pExTra02 | Fruitloop *52* | Toxic | - |
| 1 | pExTra-Hammy47 | Hammy *47* replicate 1 | Non-toxic | - |
| 2 | pExTra-Hammy47 | Hammy *47* replicate 2 | Non-toxic | - |
| 3 | pExTra-Hammy47 | Hammy *47* replicate 3 | Non-toxic | - |

*Key: NG (no growth) - (no pink color) +(faint pink color) ++(obvious pink color) +++ (dark pink color)

### Gene *48*; Score *0*

Images taken after 3 days at 37 ºC

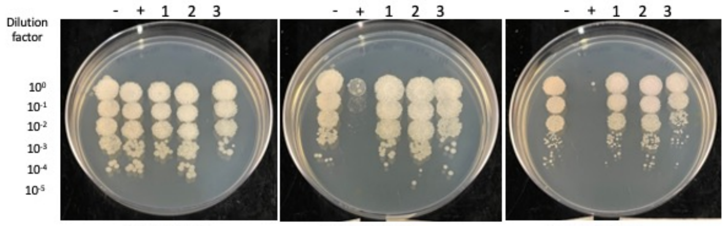


0 ng/ml aTc | 10 ng/ml aTc | 100 ng/ml aTc

| Lane | Plasmid name | Gene name, replicate | Toxic/Non-toxic | Colony color on 100 ng/ml aTc plate* |
|---|---|---|---|---|
| - Non-toxic control | pExTra03 | Fruitloop *52* mutant | Non-toxic | - |
| + Toxic control | pExTra02 | Fruitloop *52* | Toxic | -/NG |
| 1 | pExTra-Hammy48 | Hammy *48* replicate 1 | Non-toxic | - |
| 2 | pExTra-Hammy48 | Hammy *48* replicate 2 | Non-toxic | - |
| 3 | pExTra-Hammy48 | Hammy *48* replicate 3 | Non-toxic | - |

*Key: NG (no growth) - (no pink color) +(faint pink color) ++(obvious pink color) +++ (dark pink color)

### Gene *49*; Score *0*

Images taken after 4 days at 37 ºC

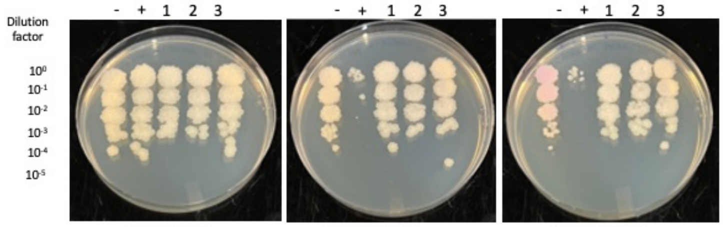


0 ng/ml aTc | 10 ng/ml aTc | 100 ng/ml aTc

| Lane | Plasmid name | Gene name, replicate | Toxic/Non-toxic | Colony color on 100 ng/ml aTc plate* |
|---|---|---|---|---|
| - Non-toxic control | pExTra03 | Fruitloop *52* mutant | Non-toxic | ++ |
| + Toxic control | pExTra02 | Fruitloop *52* | Toxic | - |
| 1 | pExTra-Hammy49 | Hammy *49* replicate 1 | Non-toxic | - |
| 2 | pExTra-Hammy49 | Hammy *49* replicate 2 | Non-toxic | - |
| 3 | pExTra-Hammy49 | Hammy 49 replicate 3 | Non-toxic | - |

*Key: NG (no growth) - (no pink color) +(faint pink color) ++(obvious pink color) +++ (dark pink color)

### Gene *50*; Score *3*

Images taken after 4 days at 37 ºC

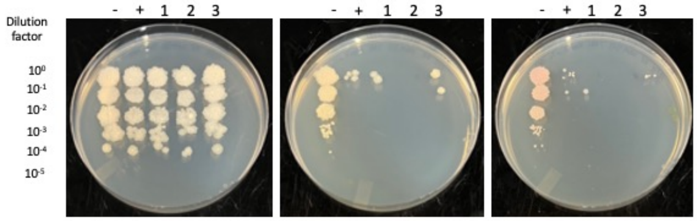


0 ng/ml aTc | 10 ng/ml aTc | 100 ng/ml aTc

| Lane | Plasmid name | Gene name, replicate | Toxic/Non-toxic | Colony color on 100 ng/ml aTc plate* |
|---|---|---|---|---|
| - Non-toxic control | pExTra03 | Fruitloop *52* mutant | Non-toxic | + |
| + Toxic control | pExTra02 | Fruitloop *52* | Toxic | -/NG |
| 1 | pExTra-Hammy50 | Hammy *50* replicate 1 | Toxic | NG |
| 2 | pExTra-Hammy50 | Hammy *50* replicate 2 | Toxic | NG |
| 3 | pExTra-Hammy50 | Hammy *50* replicate 3 | Toxic | -/NG |

*Key: NG (no growth) - (no pink color) +(faint pink color) ++(obvious pink color) +++ (dark pink color)

Images taken after 3 days at 37 ºC

Gene *51*; Score *3*

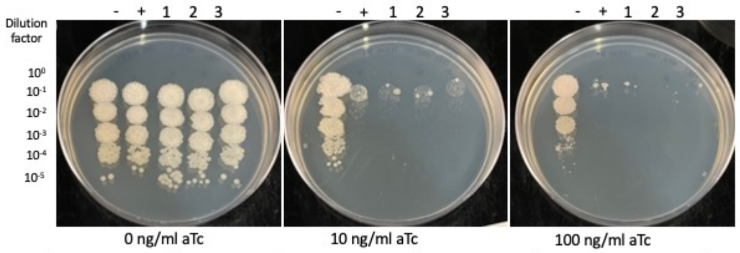


| Lane | Plasmid name | Gene name, replicate | Toxic/Non-toxic | Colony color on 100 ng/ml aTc plate* |
|---|---|---|---|---|
| - Non-toxic control | pExTra03 | Fruitloop *52* mutant | Non-toxic | + |
| + Toxic control | pExTra02 | Fruitloop *52* | Toxic | - |
| 1 | pExTra-Hammy51 | Hammy *51* replicate 1 | Toxic | -/NG |
| 2 | pExTra-Hammy51 | Hammy *51* replicate 2 | Toxic | NG |
| 3 | pExTra-Hammy51 | Hammy *51* replicate 3 | Toxic | NG |

*Key: NG (no growth) - (no pink color) +(faint pink color) ++(obvious pink color) +++ (dark pink color)

Images taken after 4 days at 37 ºC

Gene *52*; Score *0*

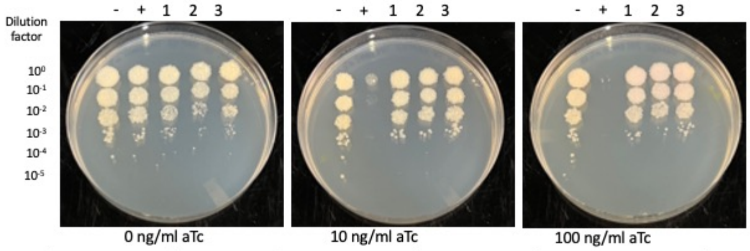


| Lane | Plasmid name | Gene name, replicate | Toxic/Non-toxic | Colony color on 100 ng/ml aTc plate* |
|---|---|---|---|---|
| - Non-toxic control | pExTra03 | Fruitloop *52* mutant | Non-toxic | - |
| + Toxic control | pExTra02 | Fruitloop *52* | Toxic | - |
| 1 | pExTra-Hammy52 | Hammy *52* replicate 1 | Non-toxic | + |
| 2 | pExTra-Hammy52 | Hammy *52* replicate 2 | Non-toxic | + |
| 3 | pExTra-Hammy52 | Hammy *52* replicate 3 | Non-toxic | + |

*Key: NG (no growth) - (no pink color) +(faint pink color) ++(obvious pink color) +++ (dark pink color)

Images taken after 4 days at 37 ºC

Gene *53*; Score *1*

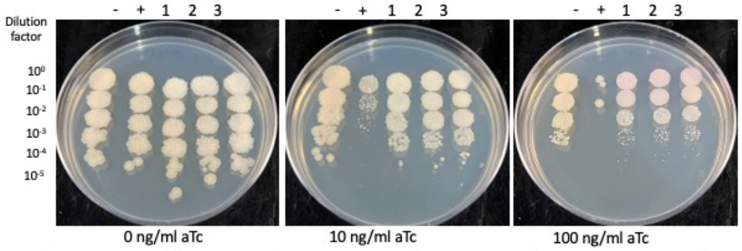


| Lane | Plasmid name | Gene name, replicate | Toxic/Non-toxic | Colony color on 100 ng/ml aTc plate* |
|---|---|---|---|---|
| - Non-toxic control | pExTra03 | Fruitloop *52* mutant | Non-toxic | + |
| + Toxic control | pExTra02 | Fruitloop *52* | Toxic | - |
| 1 | pExTra-Hammy53 | Hammy *53* replicate 1 | Toxic | ++ |
| 2 | pExTra-Hammy53 | Hammy *53* replicate 2 | Toxic | ++ |
| 3 | pExTra-Hammy53 | Hammy *53* replicate 3 | Toxic | ++ |

*Key: NG (no growth) - (no pink color) +(faint pink color) ++(obvious pink color) +++ (dark pink color)

Images taken after 3 days at 37 ºC

Gene *54*; Score *3*

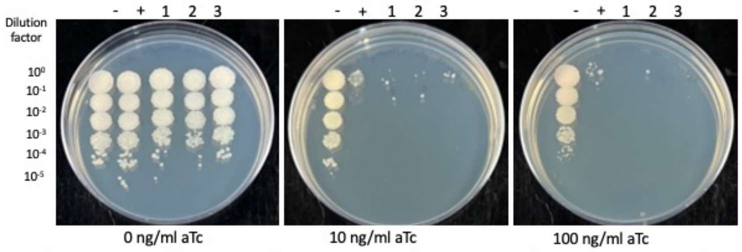


| Lane | Plasmid name | Gene name, replicate | Toxic/Non-toxic | Colony color on 100 ng/ml aTc plate* |
|---|---|---|---|---|
| - Non-toxic control | pExTra03 | Fruitloop *52* mutant | Non-toxic | + |
| + Toxic control | pExTra02 | Fruitloop *52* | Toxic | - |
| 1 | pExTra-Hammy54 | Hammy *54* replicate 1 | Toxic | NG |
| 2 | pExTra-Hammy54 | Hammy *54* replicate 2 | Toxic | NG |
| 3 | pExTra-Hammy54 | Hammy *54* replicate 3 | Toxic | NG |

*Key: NG (no growth) - (no pink color) +(faint pink color) ++(obvious pink color) +++ (dark pink color)

Images taken after 4 days at 37 ºC

Gene *55*; Score *0*

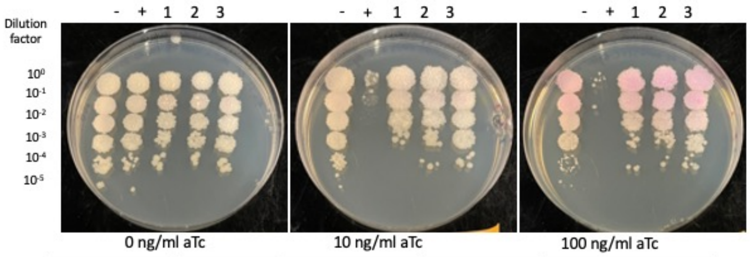


| Lane | Plasmid name | Gene name, replicate | Toxic/Non-toxic | Colony color on 100 ng/ml aTc plate* |
|---|---|---|---|---|
| - Non-toxic control | pExTra03 | Fruitloop *52* mutant | Non-toxic | ++ |
| + Toxic control | pExTra02 | Fruitloop *52* | Toxic | - |
| 1 | pExTra-Hammy55 | Hammy *55* replicate 1 | Non-toxic | ++ |
| 2 | pExTra-Hammy55 | Hammy *55* replicate 2 | Non-toxic | ++ |
| 3 | pExTra-Hammy55 | Hammy *55* replicate 3 | Non-toxic | ++ |

*Key: NG (no growth) - (no pink color) +(faint pink color) ++(obvious pink color) +++ (dark pink color)

Images taken after 4 days at 37 ºC

Gene *56*; Score *3*

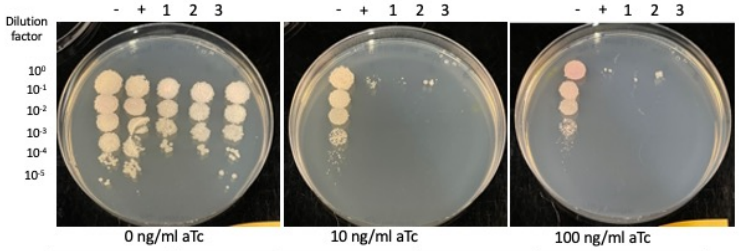


| Lane | Plasmid name | Gene name, replicate | Toxic/Non-toxic | Colony color on 100 ng/ml aTc plate* |
|---|---|---|---|---|
| - Non-toxic control | pExTra03 | Fruitloop *52* mutant | Non-toxic | + |
| + Toxic control | pExTra02 | Fruitloop *52* | Toxic | -/NG |
| 1 | pExTra-Hammy56 | Hammy *56* replicate 1 | Toxic | -/NG |
| 2 | pExTra-Hammy56 | Hammy *56* replicate 2 | Toxic | -/NG |
| 3 | pExTra-Hammy56 | Hammy *56* replicate 3 | Toxic | NG |

*Key: NG (no growth) - (no pink color) +(faint pink color) ++(obvious pink color) +++ (dark pink color)

Images taken after 4 days at 37 ºC

Gene *57*; Score *0*

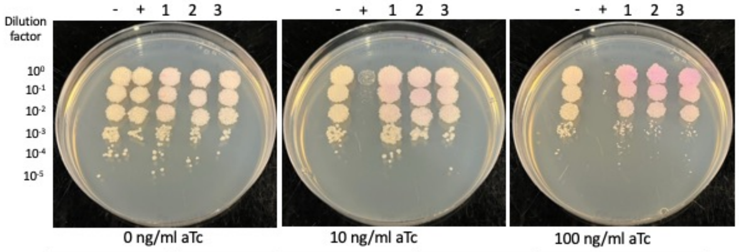


| Lane | Plasmid name | Gene name, replicate | Toxic/Non-toxic | Colony color on 100 ng/ml aTc plate* |
|---|---|---|---|---|
| - Non-toxic control | pExTra03 | Fruitloop *52* mutant | Non-toxic | + |
| + Toxic control | pExTra02 | Fruitloop *52* | Toxic | - |
| 1 | pExTra-Hammy57 | Hammy *57* replicate 1 | Non-toxic | ++ |
| 2 | pExTra-Hammy57 | Hammy *57* replicate 2 | Non-toxic | ++ |
| 3 | pExTra-Hammy57 | Hammy *57* replicate 3 | Non-toxic | ++ |

*Key: NG (no growth) - (no pink color) +(faint pink color) ++(obvious pink color) +++ (dark pink color)

Images taken after 3 days at 37 ºC

Gene *58*; Score *3*

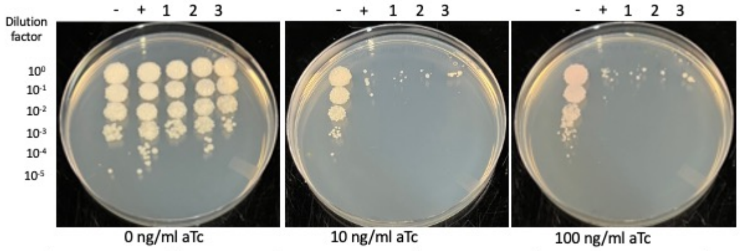


| Lane | Plasmid name | Gene name, replicate | Toxic/Non-toxic | Colony color on 100 ng/ml aTc plate* |
|---|---|---|---|---|
| - Non-toxic control | pExTra03 | Fruitloop *52* mutant | Non-toxic | + |
| + Toxic control | pExTra02 | Fruitloop *52* | Toxic | -/NG |
| 1 | pExTra-Hammy58 | Hammy *58* replicate 1 | Toxic | -/NG |
| 2 | pExTra-Hammy58 | Hammy *58* replicate 2 | Toxic | -/NG |
| 3 | pExTra-Hammy58 | Hammy *58* replicate 3 | Toxic | -/NG |

*Key: NG (no growth) - (no pink color) +(faint pink color) ++(obvious pink color) +++ (dark pink color)

Images taken after 6 days at 37 ºC

### Gene *59*; Score *0*

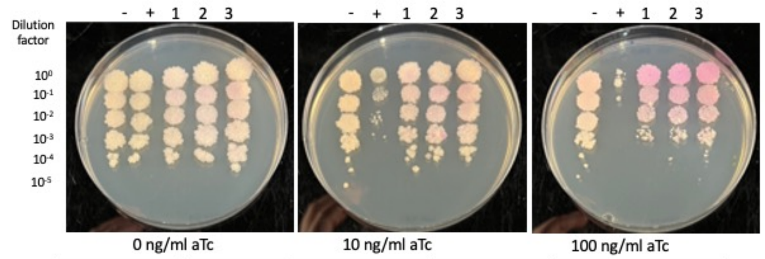


| Lane | Plasmid name | Gene name, replicate | Toxic/Non-toxic | Colony color on 100 ng/ml aTc plate* |
|---|---|---|---|---|
| - Non-toxic control | pExTra03 | Fruitloop *52* mutant | Non-toxic | + |
| + Toxic control | pExTra02 | Fruitloop *52* | Toxic | - |
| 1 | pExTra-Hammy59 | Hammy *59* replicate 1 | Non-toxic | +++ |
| 2 | pExTra-Hammy59 | Hammy *59* replicate 2 | Non-toxic | +++ |
| 3 | pExTra-Hammy59 | Hammy *59* replicate 3 | Non-toxic | +++ |

*Key: NG (no growth) - (no pink color) +(faint pink color) ++(obvious pink color) +++ (dark pink color)

Images taken after 3 days at 37 ºC

### Gene *60*; Score *1*

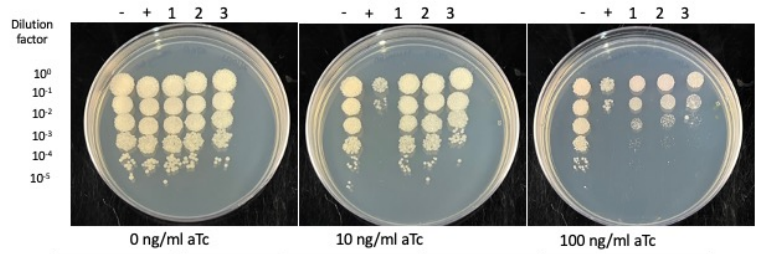


| Lane | Plasmid name | Gene name, replicate | Toxic/Non-toxic | Colony color on 100 ng/ml aTc plate* |
|---|---|---|---|---|
| - Non-toxic control | pExTra03 | Fruitloop *52* mutant | Non-toxic | - |
| + Toxic control | pExTra02 | Fruitloop *52* | Toxic | - |
| 1 | pExTra-Hammy60 | Hammy *60* replicate 1 | Toxic | - |
| 2 | pExTra-Hammy60 | Hammy *60* replicate 2 | Toxic | - |
| 3 | pExTra-Hammy60 | Hammy *60* replicate 3 | Toxic | - |

*Key: NG (no growth) - (no pink color) +(faint pink color) ++(obvious pink color) +++ (dark pink color)

Images taken after 3 days at 37 ºC

### Gene *61*; Score *1*

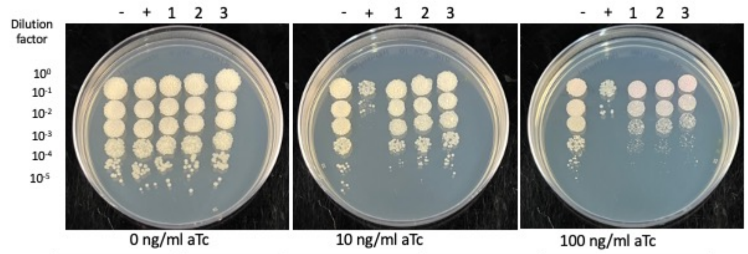


| Lane | Plasmid name | Gene name, replicate | Toxic/Non-toxic | Colony color on 100 ng/ml aTc plate* |
|---|---|---|---|---|
| - Non-toxic control | pExTra03 | Fruitloop *52* mutant | Non-toxic | - |
| + Toxic control | pExTra02 | Fruitloop *52* | Toxic | - |
| 1 | pExTra-Hammy61 | Hammy *61* replicate 1 | Toxic | - |
| 2 | pExTra-Hammy61 | Hammy *61* replicate 2 | Toxic | - |
| 3 | pExTra-Hammy61 | Hammy *61* replicate 3 | Toxic | - |

*Key: NG (no growth) - (no pink color) +(faint pink color) ++(obvious pink color) +++ (dark pink color)

Images taken after 3 days at 37 ºC

### Gene *62*; Score *0*

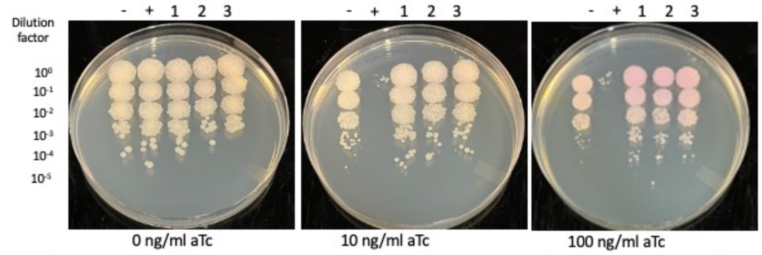


| Lane | Plasmid name | Gene name, replicate | Toxic/Non-toxic | Colony color on 100 ng/ml aTc plate* |
|---|---|---|---|---|
| - Non-toxic control | pExTra03 | Fruitloop *52* mutant | Non-toxic | + |
| + Toxic control | pExTra02 | Fruitloop *52* | Toxic | - |
| 1 | pExTra-Hammy62 | Hammy *62* replicate 1 | Non-toxic | ++ |
| 2 | pExTra-Hammy62 | Hammy *62* replicate 2 | Non-toxic | ++ |
| 3 | pExTra-Hammy62 | Hammy *62* replicate 3 | Non-toxic | ++ |

*Key: NG (no growth) - (no pink color) +(faint pink color) ++(obvious pink color) +++ (dark pink color)

Images taken after 3 days at 37 ºC

### Gene *63*; Score *1*

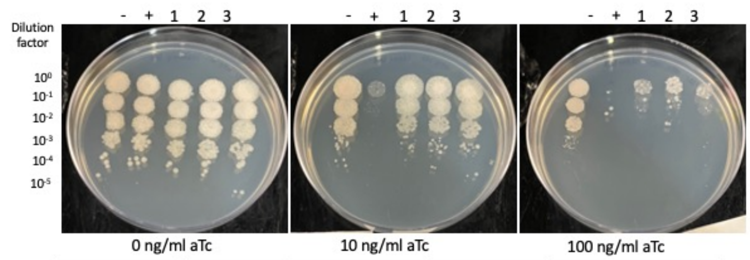


| Lane | Plasmid name | Gene name, replicate | Toxic/Non-toxic | Colony color on 100 ng/ml aTc plate* |
|---|---|---|---|---|
| - Non-toxic control | pExTra03 | Fruitloop *52* mutant | Non-toxic | - |
| + Toxic control | pExTra02 | Fruitloop *52* | Toxic | - |
| 1 | pExTra-Hammy63 | Hammy *63* replicate 1 | Toxic | - |
| 2 | pExTra-Hammy63 | Hammy *63* replicate 2 | Toxic | - |
| 3 | pExTra-Hammy63 | Hammy *63* replicate 3 | Toxic | - |

*Key: NG (no growth) - (no pink color) +(faint pink color) ++(obvious pink color) +++ (dark pink color)

Images taken after 4 days at 37 ºC

### Gene *64*; Score *0*

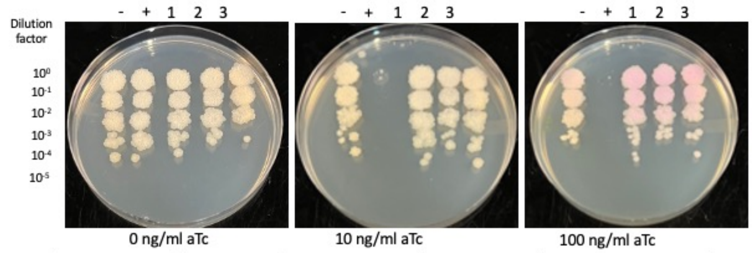


| Lane | Plasmid name | Gene name, replicate | Toxic/Non-toxic | Colony color on 100 ng/ml aTc plate* |
|---|---|---|---|---|
| - Non-toxic control | pExTra03 | Fruitloop *52* mutant | Non-toxic | + |
| + Toxic control | pExTra02 | Fruitloop *52* | Toxic | - |
| 1 | pExTra-Hammy64 | Hammy *64* replicate 1 | Non-toxic | ++ |
| 2 | pExTra-Hammy64 | Hammy *64* replicate 2 | Non-toxic | ++ |
| 3 | pExTra-Hammy64 | Hammy *64* replicate 3 | Non-toxic | ++ |

*Key: NG (no growth) - (no pink color) +(faint pink color) ++(obvious pink color) +++ (dark pink color)

Images taken after 3 days at 37 ºC

### Gene *65*; Score *0*

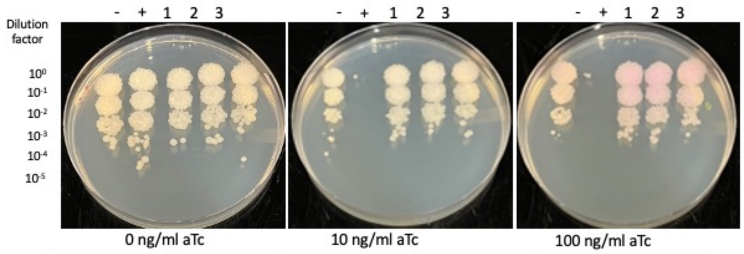


| Lane | Plasmid name | Gene name, replicate | Toxic/Non-toxic | Colony color on 100 ng/ml aTc plate* |
|---|---|---|---|---|
| - Non-toxic control | pExTra03 | Fruitloop *52* mutant | Non-toxic | + |
| + Toxic control | pExTra02 | Fruitloop *52* | Toxic | - |
| 1 | pExTra-Hammy65 | Hammy *65* replicate 1 | Non-toxic | ++ |
| 2 | pExTra-Hammy65 | Hammy *65* replicate 2 | Non-toxic | ++ |
| 3 | pExTra-Hammy65 | Hammy *65* replicate 3 | Non-toxic | ++ |

*Key: NG (no growth) - (no pink color) +(faint pink color) ++(obvious pink color) +++ (dark pink color)

Images taken after 3 days at 37 ºC

### Gene *66*; Score *0*

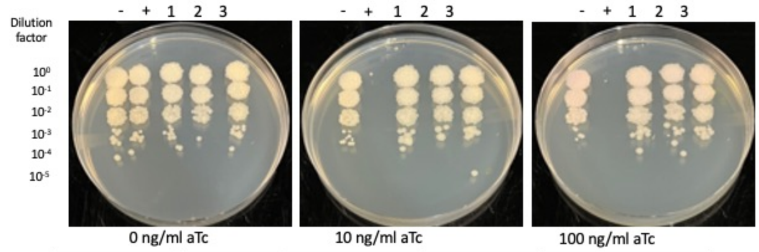


| Lane | Plasmid name | Gene name, replicate | Toxic/Non-toxic | Colony color on 100 ng/ml aTc plate* |
|---|---|---|---|---|
| - Non-toxic control | pExTra03 | Fruitloop *52* mutant | Non-toxic | + |
| + Toxic control | pExTra02 | Fruitloop *52* | Toxic | - |
| 1 | pExTra-Hammy66 | Hammy *66* replicate 1 | Non-toxic | + |
| 2 | pExTra-Hammy66 | Hammy *66* replicate 2 | Non-toxic | + |
| 3 | pExTra-Hammy66 | Hammy *66* replicate 3 | Non-toxic | + |

*Key: NG (no growth) - (no pink color) +(faint pink color) ++(obvious pink color) +++ (dark pink color)

Images taken after 4 days at 37 ºC

### Gene *67*; Score *1*

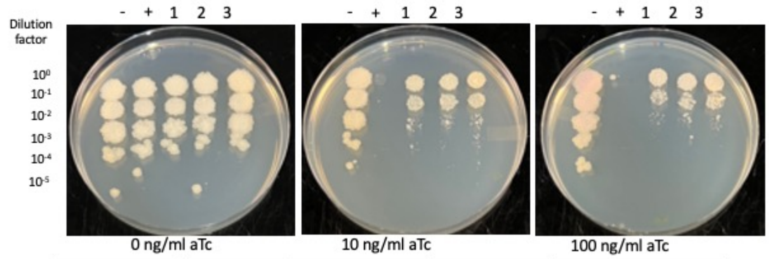


| Lane | Plasmid name | Gene name, replicate | Toxic/Non-toxic | Colony color on 100 ng/ml aTc plate* |
|---|---|---|---|---|
| - Non-toxic control | pExTra03 | Fruitloop *52* mutant | Non-toxic | + |
| + Toxic control | pExTra02 | Fruitloop *52* | Toxic | - |
| 1 | pExTra-Hammy67 | Hammy *67* replicate 1 | Toxic | - |
| 2 | pExTra-Hammy67 | Hammy *67* replicate 2 | Toxic | - |
| 3 | pExTra-Hammy67 | Hammy *67* replicate 3 | Toxic | - |

*Key: NG (no growth) - (no pink color) +(faint pink color) ++(obvious pink color) +++ (dark pink color)

Images taken after 3 days at 37 ºC

### Gene *68*; Score *3*

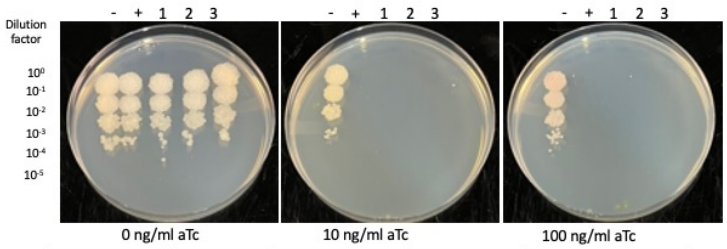


| Lane | Plasmid name | Gene name, replicate | Toxic/Non-toxic | Colony color on 100 ng/ml aTc plate* |
|---|---|---|---|---|
| - Non-toxic control | pExTra03 | Fruitloop *52* mutant | Non-toxic | + |
| + Toxic control | pExTra02 | Fruitloop *52* | Toxic | NG |
| 1 | pExTra-Hammy68 | Hammy *68* replicate 1 | Toxic | NG |
| 2 | pExTra-Hammy68 | Hammy *68* replicate 2 | Toxic | NG |
| 3 | pExTra-Hammy68 | Hammy *68* replicate 3 | Toxic | NG |

*Key: NG (no growth) - (no pink color) +(faint pink color) ++(obvious pink color) +++ (dark pink color)

Images taken after 3 days at 37 ºC

### Gene *69*; Score *3*

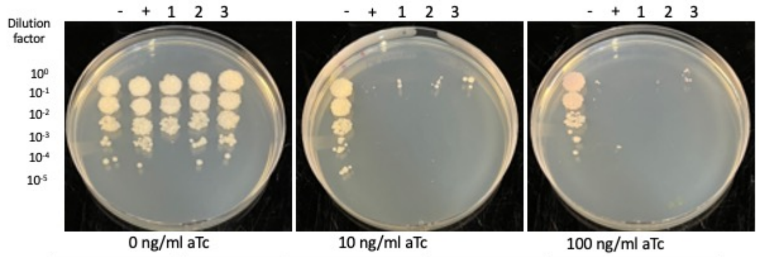


| Lane | Plasmid name | Gene name, replicate | Toxic/Non-toxic | Colony color on 100 ng/ml aTc plate* |
|---|---|---|---|---|
| - Non-toxic control | pExTra03 | Fruitloop *52* mutant | Non-toxic | + |
| + Toxic control | pExTra02 | Fruitloop *52* | Toxic | -/NG |
| 1 | pExTra-Hammy69 | Hammy *69* replicate 1 | Toxic | -/NG |
| 2 | pExTra-Hammy69 | Hammy *69* replicate 2 | Toxic | -/NG |
| 3 | pExTra-Hammy69 | Hammy *69* replicate 3 | Toxic | -/NG |

*Key: NG (no growth) - (no pink color) +(faint pink color) ++(obvious pink color) +++ (dark pink color)

Images taken after 4 days at 37 ºC

### Gene *70*; Score *0*

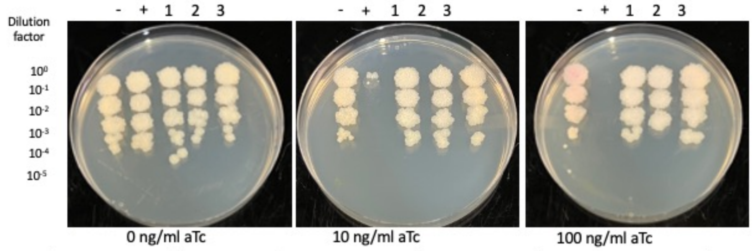


| Lane | Plasmid name | Gene name, replicate | Toxic/Non-toxic | Colony color on 100 ng/ml aTc plate* |
|---|---|---|---|---|
| - Non-toxic control | pExTra03 | Fruitloop *52* mutant | Non-toxic | + |
| + Toxic control | pExTra02 | Fruitloop *52* | Toxic | NG |
| 1 | pExTra-Hammy70 | Hammy *70* replicate 1 | Non-toxic | - |
| 2 | pExTra-Hammy70 | Hammy *70* replicate 2 | Non-toxic | - |
| 3 | pExTra-Hammy70 | Hammy *70* replicate 3 | Non-toxic | - |

*Key: NG (no growth) - (no pink color) +(faint pink color) ++(obvious pink color) +++ (dark pink color)

Images taken after 5 days at 37 ºC

### Gene *71*; Score *0*

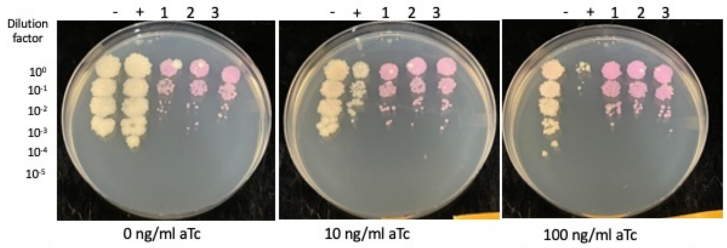


| Lane | Plasmid name | Gene name, replicate | Toxic/Non-toxic | Colony color on 100 ng/ml aTc plate* |
|---|---|---|---|---|
| - Non-toxic control | pExTra03 | Fruitloop *52* mutant | Non-toxic | + |
| + Toxic control | pExTra02 | Fruitloop *52* | Toxic | - |
| 1 | pExTra-Hammy71 | Hammy *71* replicate 1 | Non-toxic | +++ |
| 2 | pExTra-Hammy71 | Hammy *71* replicate 2 | Non-toxic | +++ |
| 3 | pExTra-Hammy71 | Hammy *71* replicate 3 | Non-toxic | +++ |

*Key: NG (no growth) - (no pink color) +(faint pink color) ++(obvious pink color) +++ (dark pink color)

Images taken after 4 days at 37 ºC

### Gene *72*; Score *0*

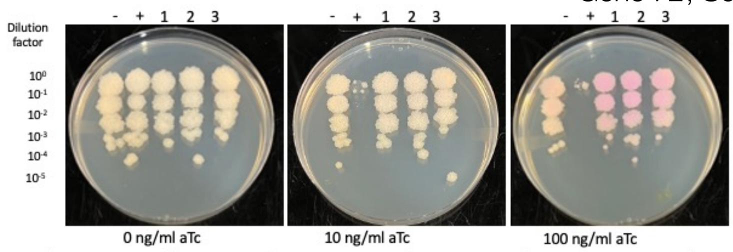


| Lane | Plasmid name | Gene name, replicate | Toxic/Non-toxic | Colony color on 100 ng/ml aTc plate* |
|---|---|---|---|---|
| - Non-toxic control | pExTra03 | Fruitloop *52* mutant | Non-toxic | + |
| + Toxic control | pExTra02 | Fruitloop *52* | Toxic | + |
| 1 | pExTra-Hammy72 | Hammy *72* replicate 1 | Non-toxic | ++ |
| 2 | pExTra-Hammy72 | Hammy *72* replicate 2 | Non-toxic | ++ |
| 3 | pExTra-Hammy72 | Hammy *72* replicate 3 | Non-toxic | ++ |

*Key: NG (no growth) - (no pink color) +(faint pink color) ++(obvious pink color) +++ (dark pink color)

Images taken after 4 days at 37 ºC

### Gene *73*; Score *0*

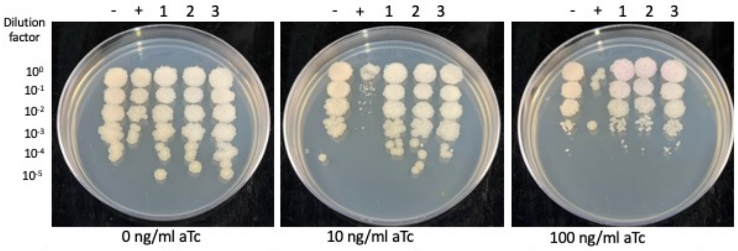


| Lane | Plasmid name | Gene name, replicate | Toxic/Non-toxic | Colony color on 100 ng/ml aTc plate* |
|---|---|---|---|---|
| - Non-toxic control | pExTra03 | Fruitloop *52* mutant | Non-toxic | + |
| + Toxic control | pExTra02 | Fruitloop *52* | Toxic | - |
| 1 | pExTra-Hammy73 | Hammy *73* replicate 1 | Non-toxic | + |
| 2 | pExTra-Hammy73 | Hammy *73* replicate 2 | Non-toxic | + |
| 3 | pExTra-Hammy73 | Hammy *73* replicate 3 | Non-toxic | + |

*Key: NG (no growth) - (no pink color) +(faint pink color) ++(obvious pink color) +++ (dark pink color)

Images taken after 4 days at 37 ºC

### Gene *74*; Score *0*

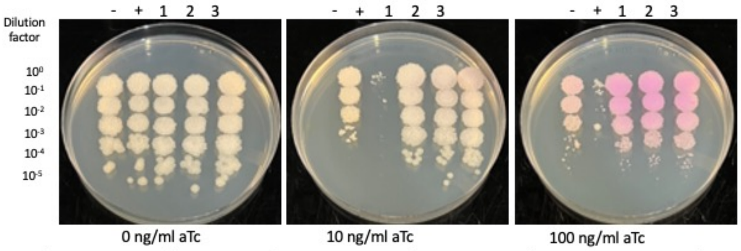


| Lane | Plasmid name | Gene name, replicate | Toxic/Non-toxic | Colony color on 100 ng/ml aTc plate* |
|---|---|---|---|---|
| - Non-toxic control | pExTra03 | Fruitloop *52* mutant | Non-toxic | ++ |
| + Toxic control | pExTra02 | Fruitloop *52* | Toxic | - |
| 1 | pExTra-Hammy74 | Hammy *74* replicate 1 | Non-toxic | +++ |
| 2 | pExTra-Hammy74 | Hammy *74* replicate 2 | Non-toxic | +++ |
| 3 | pExTra-Hammy74 | Hammy *74* replicate 3 | Non-toxic | +++ |

*Key: NG (no growth) - (no pink color) +(faint pink color) ++(obvious pink color) +++ (dark pink color)

### Gene *75*; Score *0*

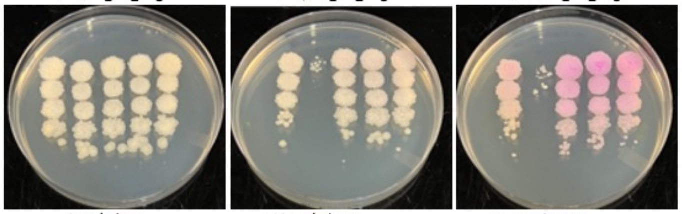


| Lane | Plasmid name | Gene name, replicate | Toxic/Non-toxic | Colony color on 100 ng/ml aTc plate* |
|---|---|---|---|---|
| - Non-toxic control | pExTra03 | Fruitloop *52* mutant | Non-toxic | ++ |
| + Toxic control | pExTra02 | Fruitloop *52* | Toxic | - |
| 1 | pExTra-Hammy75 | Hammy *75* replicate 1 | Non-toxic | +++ |
| 2 | pExTra-Hammy75 | Hammy *75* replicate 2 | Non-toxic | +++ |
| 3 | pExTra-Hammy75 | Hammy *75* replicate 3 | Non-toxic | +++ |

*Key: NG (no growth) - (no pink color) +(faint pink color) ++(obvious pink color) +++ (dark pink color)

### Gene *76*; Score *0*

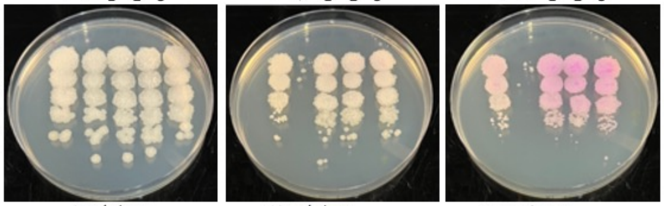


| Lane | Plasmid name | Gene name, replicate | Toxic/Non-toxic | Colony color on 100 ng/ml aTc plate* |
|---|---|---|---|---|
| - Non-toxic control | pExTra03 | Fruitloop *52* mutant | Non-toxic | + |
| + Toxic control | pExTra02 | Fruitloop *52* | Toxic | - |
| 1 | pExTra-Hammy76 | Hammy *76* replicate 1 | Non-toxic | +++ |
| 2 | pExTra-Hammy76 | Hammy *76* replicate 2 | Non-toxic | +++ |
| 3 | pExTra-Hammy76 | Hammy *76* replicate 3 | Non-toxic | +++ |

*Key: NG (no growth) - (no pink color) +(faint pink color) ++(obvious pink color) +++ (dark pink color)

### Gene *77*; Score *3*

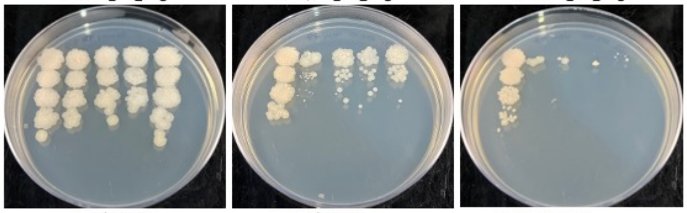


| Lane | Plasmid name | Gene name, replicate | Toxic/Non-toxic | Colony color on 100 ng/ml aTc plate* |
|---|---|---|---|---|
| - Non-toxic control | pExTra03 | Fruitloop *52* mutant | Non-toxic | + |
| + Toxic control | pExTra02 | Fruitloop *52* | Toxic | -/NG |
| 1 | pExTra-Hammy77 | Hammy *77* replicate 1 | Toxic | -/NG |
| 2 | pExTra-Hammy77 | Hammy *77* replicate 2 | Toxic | -/NG |
| 3 | pExTra-Hammy77 | Hammy *77* replicate 3 | Toxic | -/NG |

*Key: NG (no growth) - (no pink color) +(faint pink color) ++(obvious pink color) +++ (dark pink color)

### Gene *78*; Score *3*

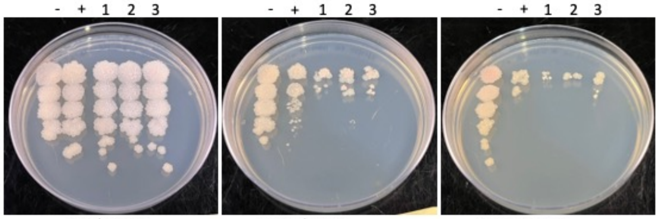


| Lane | Plasmid name | Gene name, replicate | Toxic/Non-toxic | Colony color on 100 ng/ml aTc plate* |
|---|---|---|---|---|
| - Non-toxic control | pExTra03 | Fruitloop *52* mutant | Non-toxic | + |
| + Toxic control | pExTra02 | Fruitloop *52* | Toxic | -/NG |
| 1 | pExTra-Hammy78 | Hammy *78* replicate 1 | Toxic | -/NG |
| 2 | pExTra-Hammy78 | Hammy *78* replicate 2 | Toxic | -/NG |
| 3 | pExTra-Hammy78 | Hammy *78* replicate 3 | Toxic | -/NG |

*Key: NG (no growth) - (no pink color) +(faint pink color) ++(obvious pink color) +++ (dark pink color)

### Gene *79*; Score *0*

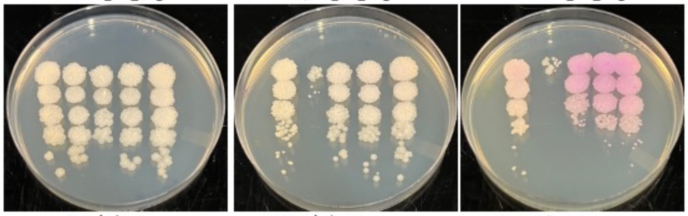


| Lane | Plasmid name | Gene name, replicate | Toxic/Non-toxic | Colony color on 100 ng/ml aTc plate* |
|---|---|---|---|---|
| - Non-toxic control | pExTra03 | Fruitloop *52* mutant | Non-toxic | + |
| + Toxic control | pExTra02 | Fruitloop *52* | Toxic | - |
| 1 | pExTra-Hammy79 | Hammy *79* replicate 1 | Non-toxic | +++ |
| 2 | pExTra-Hammy79 | Hammy *79* replicate 2 | Non-toxic | +++ |
| 3 | pExTra-Hammy79 | Hammy *79* replicate 3 | Non-toxic | +++ |

*Key: NG (no growth) - (no pink color) +(faint pink color) ++(obvious pink color) +++ (dark pink color)

### Gene *80*; Score *0*

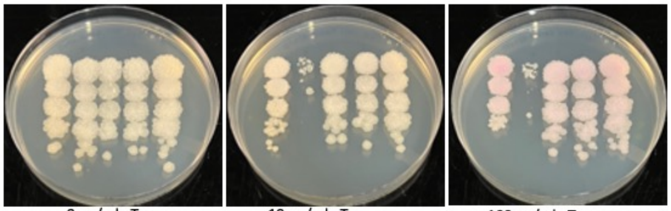


| Lane | Plasmid name | Gene name, replicate | Toxic/Non-toxic | Colony color on 100 ng/ml aTc plate* |
|---|---|---|---|---|
| - Non-toxic control | pExTra03 | Fruitloop *52* mutant | Non-toxic | + |
| + Toxic control | pExTra02 | Fruitloop *52* | Toxic | - |
| 1 | pExTra-Hammy80 | Hammy *80* replicate 1 | Non-toxic | + |
| 2 | pExTra-Hammy80 | Hammy *80* replicate 2 | Non-toxic | + |
| 3 | pExTra-Hammy80 | Hammy *80* replicate 3 | Non-toxic | + |

*Key: NG (no growth) - (no pink color) +(faint pink color) ++(obvious pink color) +++ (dark pink color)

### Gene *81*; Score *0*

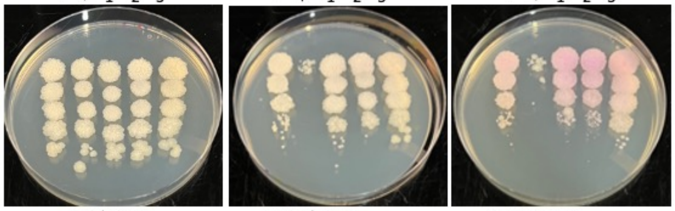


| Lane | Plasmid name | Gene name, replicate | Toxic/Non-toxic | Colony color on 100 ng/ml aTc plate* |
|---|---|---|---|---|
| - Non-toxic control | pExTra03 | Fruitloop *52* mutant | Non-toxic | + |
| + Toxic control | pExTra02 | Fruitloop *52* | Toxic | - |
| 1 | pExTra-Hammy81 | Hammy *81* replicate 1 | Non-toxic | ++ |
| 2 | pExTra-Hammy81 | Hammy *81* replicate 2 | Non-toxic | ++ |
| 3 | pExTra-Hammy81 | Hammy *81* replicate 3 | Non-toxic | + |

*Key: NG (no growth) - (no pink color) +(faint pink color) ++(obvious pink color) +++ (dark pink color)

### Gene *82*; Score *0*

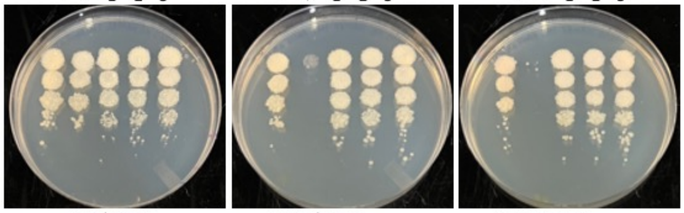


| Lane | Plasmid name | Gene name, replicate | Toxic/Non-toxic | Colony color on 100 ng/ml aTc plate* |
|---|---|---|---|---|
| - Non-toxic control | pExTra03 | Fruitloop *52* mutant | Non-toxic | - |
| + Toxic control | pExTra02 | Fruitloop *52* | Toxic | - |
| 1 | pExTra-Hammy82 | Hammy *82* replicate 1 | Non-toxic | - |
| 2 | pExTra-Hammy82 | Hammy *82* replicate 2 | Non-toxic | - |
| 3 | pExTra-Hammy82 | Hammy *82* replicate 3 | Non-toxic | - |

*Key: NG (no growth) - (no pink color) +(faint pink color) ++(obvious pink color) +++ (dark pink color)

Images taken after 3 days at 37 ºC

### Gene *83*; Score *0*

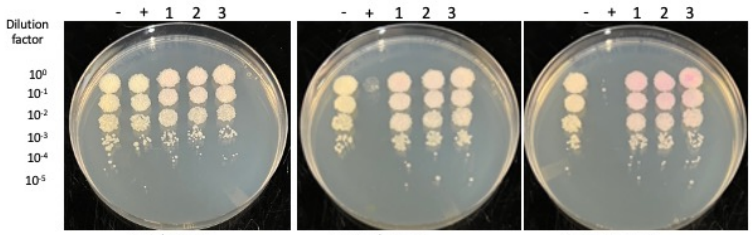


0 ng/ml aTc | 10 ng/ml aTc | 100 ng/ml aTc

| Lane | Plasmid name | Gene name, replicate | Toxic/Non-toxic | Colony color on 100 ng/ml aTc plate* |
|---|---|---|---|---|
| - Non-toxic control | pExTra03 | Fruitloop *52* mutant | Non-toxic | - |
| + Toxic control | pExTra02 | Fruitloop *52* | Toxic | - |
| 1 | pExTra-Hammy83 | Hammy *83* replicate 1 | Non-toxic | ++ |
| 2 | pExTra-Hammy83 | Hammy *83* replicate 2 | Non-toxic | ++ |
| 3 | pExTra-Hammy83 | Hammy *83* replicate 3 | Non-toxic | ++ |

*Key: NG (no growth) - (no pink color) +(faint pink color) ++(obvious pink color) +++ (dark pink color)

Images taken after 5 days at 37 ºC

### Gene *84*; Score *1*

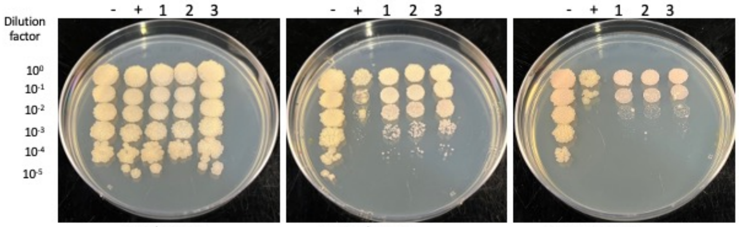


0 ng/ml aTc | 10 ng/ml aTc | 100 ng/ml aTc

| Lane | Plasmid name | Gene name, replicate | Toxic/Non-toxic | Colony color on 100 ng/ml aTc plate* |
|---|---|---|---|---|
| - Non-toxic control | pExTra03 | Fruitloop *52* mutant | Non-toxic | + |
| + Toxic control | pExTra02 | Fruitloop *52* | Toxic | - |
| 1 | pExTra-Hammy84 | Hammy *84* replicate 1 | Toxic | + |
| 2 | pExTra-Hammy84 | Hammy *84* replicate 2 | Toxic | + |
| 3 | pExTra-Hammy84 | Hammy *84* replicate 3 | Toxic | + |

*Key: NG (no growth) - (no pink color) +(faint pink color) ++(obvious pink color) +++ (dark pink color)

Images taken after 3 days at 37 ºC

### Gene *85*; Score *0*

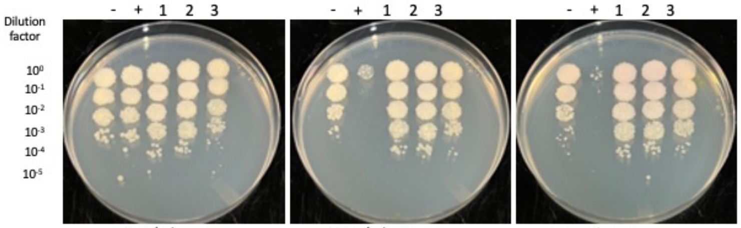


0 ng/ml aTc | 10 ng/ml aTc | 100 ng/ml aTc

| Lane | Plasmid name | Gene name, replicate | Toxic/Non-toxic | Colony color on 100 ng/ml aTc plate* |
|---|---|---|---|---|
| - Non-toxic control | pExTra03 | Fruitloop *52* mutant | Non-toxic | + |
| + Toxic control | pExTra02 | Fruitloop *52* | Toxic | - |
| 1 | pExTra-Hammy85 | Hammy *85* replicate 1 | Non-toxic | + |
| 2 | pExTra-Hammy85 | Hammy *85* replicate 2 | Non-toxic | + |
| 3 | pExTra-Hammy85 | Hammy *85* replicate 3 | Non-toxic | + |

*Key: NG (no growth) - (no pink color) +(faint pink color) ++(obvious pink color) +++ (dark pink color)

Images taken after 3 days at 37 ºC

### Gene *86*; Score *0*

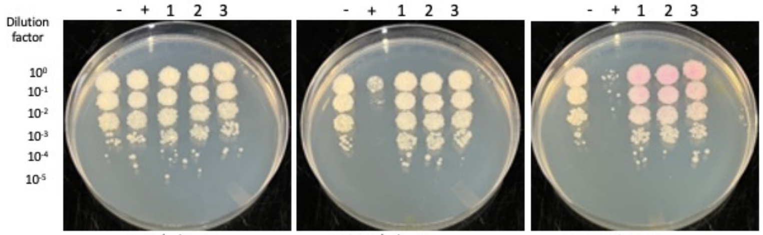


0 ng/ml aTc | 10 ng/ml aTc | 100 ng/ml aTc

| Lane | Plasmid name | Gene name, replicate | Toxic/Non-toxic | Colony color on 100 ng/ml aTc plate* |
|---|---|---|---|---|
| - Non-toxic control | pExTra03 | Fruitloop *52* mutant | Non-toxic | - |
| + Toxic control | pExTra02 | Fruitloop *52* | Toxic | - |
| 1 | pExTra-Hammy86 | Hammy *86* replicate 1 | Non-toxic | ++ |
| 2 | pExTra-Hammy86 | Hammy *86* replicate 2 | Non-toxic | ++ |
| 3 | pExTra-Hammy86 | Hammy *86* replicate 3 | Non-toxic | ++ |

*Key: NG (no growth) - (no pink color) +(faint pink color) ++(obvious pink color) +++ (dark pink color)

Images taken after 3 days at 37 ºC

### Gene *87*; Score *0*

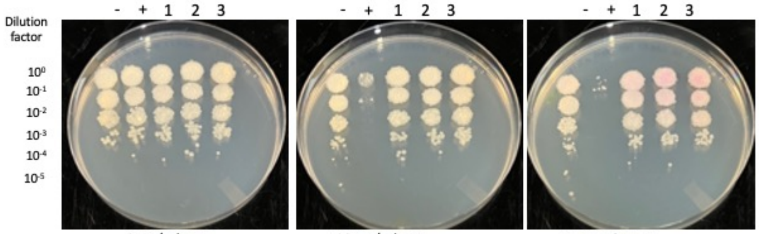


0 ng/ml aTc | 10 ng/ml aTc | 100 ng/ml aTc

| Lane | Plasmid name | Gene name, replicate | Toxic/Non-toxic | Colony color on 100 ng/ml aTc plate* |
|---|---|---|---|---|
| - Non-toxic control | pExTra03 | Fruitloop *52* mutant | Non-toxic | - |
| + Toxic control | pExTra02 | Fruitloop *52* | Toxic | - |
| 1 | pExTra-Hammy87 | Hammy *87* replicate 1 | Non-toxic | + |
| 2 | pExTra-Hammy87 | Hammy *87* replicate 2 | Non-toxic | + |
| 3 | pExTra-Hammy87 | Hammy *87* replicate 3 | Non-toxic | ++ |

*Key: NG (no growth) - (no pink color) +(faint pink color) ++(obvious pink color) +++ (dark pink color)

Images taken after 3 days at 37 ºC

### Gene *88*; Score *0*

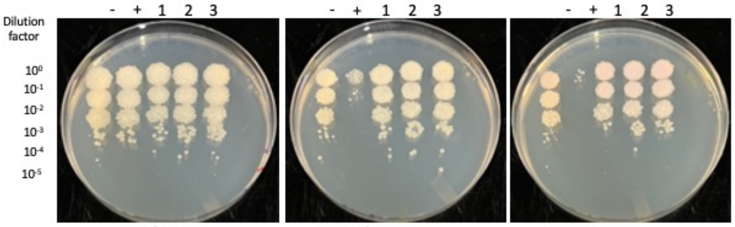


0 ng/ml aTc | 10 ng/ml aTc | 100 ng/ml aTc

| Lane | Plasmid name | Gene name, replicate | Toxic/Non-toxic | Colony color on 100 ng/ml aTc plate* |
|---|---|---|---|---|
| - Non-toxic control | pExTra03 | Fruitloop *52* mutant | Non-toxic | + |
| + Toxic control | pExTra02 | Fruitloop *52* | Toxic | - |
| 1 | pExTra-Hammy88 | Hammy *88* replicate 1 | Non-toxic | + |
| 2 | pExTra-Hammy88 | Hammy *88* replicate 2 | Non-toxic | + |
| 3 | pExTra-Hammy88 | Hammy *88* replicate 3 | Non-toxic | + |

*Key: NG (no growth) - (no pink color) +(faint pink color) ++(obvious pink color) +++ (dark pink color)

Images taken after 3 days at 37 ºC

### Gene *89*; Score *0*

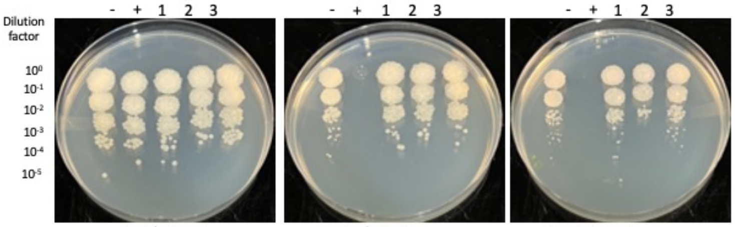


0 ng/ml aTc | 10 ng/ml aTc | 100 ng/ml aTc

| Lane | Plasmid name | Gene name, replicate | Toxic/Non-toxic | Colony color on 100 ng/ml aTc plate* |
|---|---|---|---|---|
| - Non-toxic control | pExTra03 | Fruitloop *52* mutant | Non-toxic | - |
| + Toxic control | pExTra02 | Fruitloop *52* | Toxic | - |
| 1 | pExTra-Hammy89 | Hammy *89* replicate 1 | Non-toxic | - |
| 2 | pExTra-Hammy89 | Hammy *89* replicate 2 | Non-toxic | - |
| 3 | pExTra-Hammy89 | Hammy *89* replicate 3 | Non-toxic | - |

*Key: NG (no growth) - (no pink color) +(faint pink color) ++(obvious pink color) +++ (dark pink color)

Images taken after 4 days at 37 ºC

### Gene *90*; Score *2*

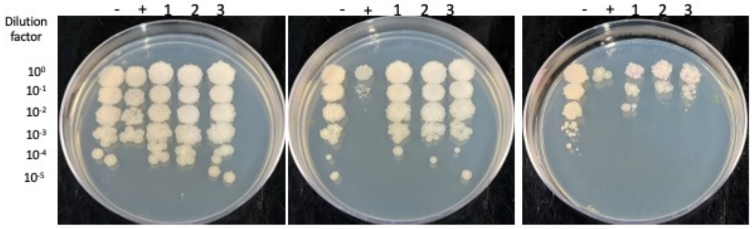


0 ng/ml aTc | 10 ng/ml aTc | 100 ng/ml aTc

| Lane | Plasmid name | Gene name, replicate | Toxic/Non-toxic | Colony color on 100 ng/ml aTc plate* |
|---|---|---|---|---|
| - Non-toxic control | pExTra03 | Fruitloop *52* mutant | Non-toxic | + |
| + Toxic control | pExTra02 | Fruitloop *52* | Toxic | - |
| 1 | pExTra-Hammy90 | Hammy *90* replicate 1 | Toxic | - |
| 2 | pExTra-Hammy90 | Hammy *90* replicate 2 | Toxic | - |
| 3 | pExTra-Hammy90 | Hammy *90* replicate 3 | Toxic | - |

*Key: NG (no growth) - (no pink color) +(faint pink color) ++(obvious pink color) +++ (dark pink color)

Images taken after 3 days at 37 ºC

### Gene *91*; Score *0*

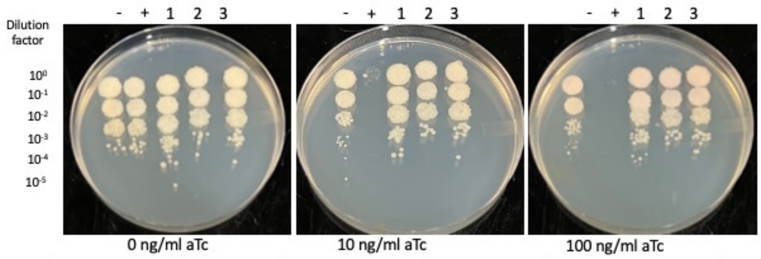


| Lane | Plasmid name | Gene name, replicate | Toxic/Non-toxic | Colony color on 100 ng/ml aTc plate* |
|---|---|---|---|---|
| - Non-toxic control | pExTra03 | Fruitloop *52* mutant | Non-toxic | + |
| + Toxic control | pExTra02 | Fruitloop *52* | Toxic | NG |
| 1 | pExTra-Hammy91 | Hammy *91* replicate 1 | Non-toxic | + |
| 2 | pExTra-Hammy91 | Hammy *91* replicate 2 | Non-toxic | + |
| 3 | pExTra-Hammy91 | Hammy *91* replicate 3 | Non-toxic | + |

*Key: NG (no growth) - (no pink color) +(faint pink color) ++(obvious pink color) +++ (dark pink color)

Images taken after 4 days at 37 ºC

### Gene *95*; Score *0*

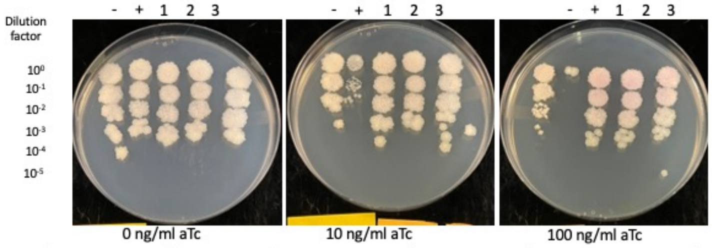


| Lane | Plasmid name | Gene name, replicate | Toxic/Non-toxic | Colony color on 100 ng/ml aTc plate* |
|---|---|---|---|---|
| - Non-toxic control | pExTra03 | Fruitloop *52* mutant | Non-toxic | + |
| + Toxic control | pExTra02 | Fruitloop *52* | Toxic | NG |
| 1 | pExTra-Hammy95 | Hammy *95* replicate 1 | Non-toxic | ++ |
| 2 | pExTra-Hammy95 | Hammy *95* replicate 2 | Non-toxic | ++ |
| 3 | pExTra-Hammy95 | Hammy *95* replicate 3 | Non-toxic | ++ |

*Key: NG (no growth) - (no pink color) +(faint pink color) ++(obvious pink color) +++ (dark pink color)

Images taken after 4 days at 37 ºC

### Gene *92*; Score *2*

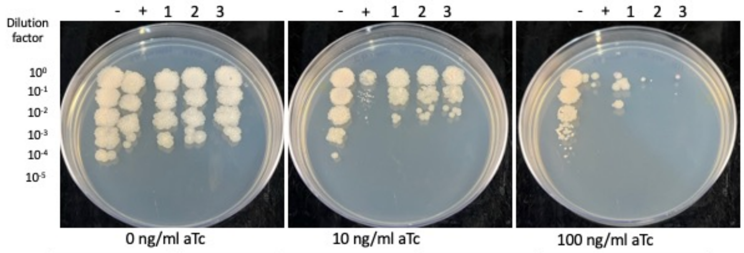


| Lane | Plasmid name | Gene name, replicate | Toxic/Non-toxic | Colony color on 100 ng/ml aTc plate* |
|---|---|---|---|---|
| - Non-toxic control | pExTra03 | Fruitloop *52* mutant | Non-toxic | + |
| + Toxic control | pExTra02 | Fruitloop *52* | Toxic | - |
| 1 | pExTra-Hammy92 | Hammy *92* replicate 1 | Toxic | -/NG |
| 2 | pExTra-Hammy92 | Hammy *92* replicate 2 | Toxic | -/NG |
| 3 | pExTra-Hammy92 | Hammy *92* replicate 3 | Toxic | NG |

*Key: NG (no growth) - (no pink color) +(faint pink color) ++(obvious pink color) +++ (dark pink color)

Images taken after 4 days at 37 ºC

### Gene *93*; Score *0*

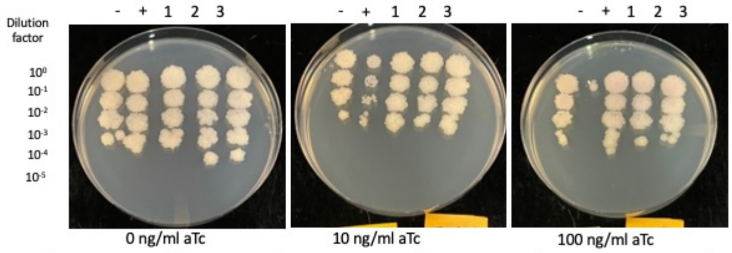


| Lane | Plasmid name | Gene name, replicate | Toxic/Non-toxic | Colony color on 100 ng/ml aTc plate* |
|---|---|---|---|---|
| - Non-toxic control | pExTra03 | Fruitloop *52* mutant | Non-toxic | + |
| + Toxic control | pExTra02 | Fruitloop *52* | Toxic | NG |
| 1 | pExTra-Hammy93 | Hammy *93* replicate 1 | Non-toxic | + |
| 2 | pExTra-Hammy93 | Hammy *93* replicate 2 | Non-toxic | + |
| 3 | pExTra-Hammy93 | Hammy *93* replicate 3 | Non-toxic | + |

*Key: NG (no growth) - (no pink color) +(faint pink color) ++(obvious pink color) +++ (dark pink color)

Images taken after 4 days at 37 ºC

### Gene *94*; Score *0*

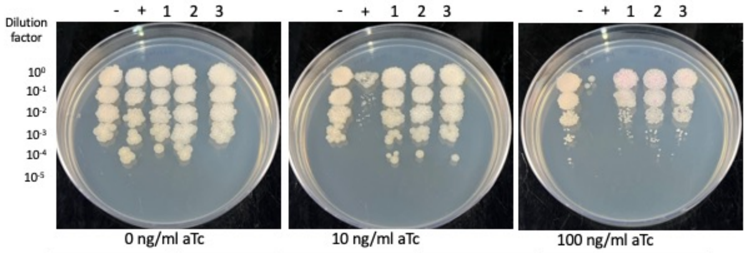


| Lane | Plasmid name | Gene name, replicate | Toxic/Non-toxic | Colony color on 100 ng/ml aTc plate* |
|---|---|---|---|---|
| - Non-toxic control | pExTra03 | Fruitloop *52* mutant | Non-toxic | + |
| + Toxic control | pExTra02 | Fruitloop *52* | Toxic | - |
| 1 | pExTra-Hammy94 | Hammy *94* replicate 1 | Non-toxic | + |
| 2 | pExTra-Hammy94 | Hammy *94* replicate 2 | Non-toxic | + |
| 3 | pExTra-Hammy94 | Hammy *94* replicate 3 | Non-toxic | + |

*Key: NG (no growth) - (no pink color) +(faint pink color) ++(obvious pink color) +++ (dark pink color)

**Supplemental Table 1: DNA oligos used in this study**

| Oligo Name | Oligo Sequence (5' to 3') |
|---|---|
| **oHammy1F** | ATGCGGAGGAATCACTTCCATATGATCGACACCTACCGCACG |
| **oHammy1R** | TGCAGGATCCGACTCGAGTGTCGACTCACTCTGCCTCGATGAAGTC |
| **oHammy2F** | ATGCGGAGGAATCACTTCCATATGACGCCGACCGTCGGGCG |
| **oHammy2R** | TGCAGGATCCGACTCGAGTGTCGACTCATGCGCGCGGGGGCCAGT |
| **oHammy3F** | ATGCGGAGGAATCACTTCCATATGAGGCCGCGGCCCGTCGG |
| **oHammy3R** | TGCAGGATCCGACTCGAGTGTCGACTCATCGGGGTGAGTCGGACAG |
| **oHammy5F** | ATGCGGAGGAATCACTTCCATATGCGTGCACCGAGCACCGTC |
| **oHammy5R** | TGCAGGATCCGACTCGAGTGTCGACTCAGCCATCGCTGCCACCGCC |
| **oHammy6F** | ATGCGGAGGAATCACTTCCATATGGCTGACGACAAGCTCGAC |
| **oHammy6R** | TGCAGGATCCGACTCGAGTGTCGACTCAGACGTCCGAACCATCGAA |
| **oHammy7F** | ATGCGGAGGAATCACTTCCATATGGTTCGGACGTCTGAGCCG |
| **oHammy7R** | TGCAGGATCCGACTCGAGTGTCGACTCACACGAACATGGCGCCACC |
| **oHammy8F** | ATGCGGAGGAATCACTTCCATATGATTCCCGCTGCCTATGAC |
| **oHammy8R** | TGCAGGATCCGACTCGAGTGTCGACTCACTGAGTCGGGCGAGCGCC |
| **oHammy9F** | ATGCGGAGGAATCACTTCCATATGACCGAGCCGACGGCGGTA |
| **oHammy9R** | TGCAGGATCCGACTCGAGTGTCGACTCATTGGTTGTAGTCCGGTGTC |
| **oHammy10F** | ATGCGGAGGAATCACTTCCATATGACGCCCGAGCAGCAGTCTT |
| **oHammy10R** | TGCAGGATCCGACTCGAGTGTCGACTCAGTCGGCGTTAGCCCGCAGG |
| **oHammy11F** | ATGCGGAGGAATCACTTCCATATGGGAGACACCGACACCGGC |
| **oHammy11R** | TGCAGGATCCGACTCGAGTGTCGACTCACTCCCCAGAGCGCAAGCG |
| **oHammy12F** | ATGCGGAGGAATCACTTCCATATGGCTGACATTTCACGCGCC |
| **oHammy12R** | TGCAGGATCCGACTCGAGTGTCGACTCAGCTACCGCCCTGCGCTGC |
| **oHammy13F** | ATGCGGAGGAATCACTTCCATATGGCACTGGCGACGACAGCG |
| **oHammy13R** | TGCAGGATCCGACTCGAGTGTCGACTCAGTAACGCTCACTGCCCA |
| **oHammy14F** | ATGCGGAGGAATCACTTCCATATGTTCCCGACGCCTCACACG |
| **oHammy14R** | TGCAGGATCCGACTCGAGTGTCGACCTATACGCCATCGTCGCCCCCCT |
| **oHammy15F** | ATGCGGAGGAATCACTTCCATATGGCGTATAGGCCCCTCGAT |
| **oHammy15R** | TGCAGGATCCGACTCGAGTGTCGACTCACAGCAGCCCACCCCTCGG |
| **oHammy16F** | ATGCGGAGGAATCACTTCCATATGACGGTCCTACTGCCTCCC |
| **oHammy16R** | TGCAGGATCCGACTCGAGTGTCGACTCAGCCAGGCCGTAGGCCGAT |
| **oHammy17F** | ATGCGGAGGAATCACTTCCATATGACCGGACCCGTTACACCG |
| **oHammy17R** | TGCAGGATCCGACTCGAGTGTCGACTCAAGCGGCGACGACCTGACCG |
| **oHammy18F** | ATGCGGAGGAATCACTTCCATATGGCAACCGCCAAGAGCAAG |
| **oHammy18R** | TGCAGGATCCGACTCGAGTGTCGACCTACCTGCCGCGCCGCCGCGC |
| **oHammy20F** | ATGCGGAGGAATCACTTCCATATGAGCGCAACGTATTACCTCAC |
| **oHammy20R** | TGCAGGATCCGACTCGAGTGTCGACTCAGGTGTAGCGCCTGCGGCTGT |
| **oHammy21F** | ATGCGGAGGAATCACTTCCATATGACCGGCACTGGTATTCACG |

| | |
|---|---|
| **oHammy21R** | TGCAGGATCCGACTCGAGTGTCGACTCAGGACCACGCCATGCGGTAG |
| **oHammy22F** | ATGCGGAGGAATCACTTCCATATGTACGTCAAGAACGGCCGCA |
| **oHammy22R** | TGCAGGATCCGACTCGAGTGTCGACCTAGAACATGTCTCCTGATCCAAAC |
| **oHammy23F** | ATGCGGAGGAATCACTTCCATATGGATCTGCCAGCCCTGCCA |
| **oHammy23R** | TGCAGGATCCGACTCGAGTGTCGACTCAGCTGTTGGCGGGCTCGTC |
| **oHammy24F** | ATGCGGAGGAATCACTTCCATATGGCCGAGGTTGAGCAGCGG |
| **oHammy24R** | TGCAGGATCCGACTCGAGTGTCGACTCAGCTCCCTTGTGGGACAACGA |
| **oHammy25F** | ATGCGGAGGAATCACTTCCATATGCCGTACACCAAGAGCTACC |
| **oHammy25R** | TGCAGGATCCGACTCGAGTGTCGACTCAGTCACCCCCCGCCTTCACC |
| **oHammy26F** | ATGCGGAGGAATCACTTCCATATGCCGTCCATGTATGACCGGC |
| **oHammy26R** | TGCAGGATCCGACTCGAGTGTCGACTCACGCCAGACTCCTATTCAGT |
| **oHammy27F** | ATGCGGAGGAATCACTTCCATATGACCGGATGGACACCAGACC |
| **oHammy27R** | TGCAGGATCCGACTCGAGTGTCGACTCACTGGTAGGCGCGGAACCATG |
| **oHammy28F** | ATGCGGAGGAATCACTTCCATATGGCTGCCACGAATCAGTTCAA |
| **oHammy28R** | TGCAGGATCCGACTCGAGTGTCGACTCAGACGGGATCGCCGTACGTGT |
| **oHammy29F** | ATGCGGAGGAATCACTTCCATATGGCTGAAAAGGTACTGCCTTAC |
| **oHammy29R** | TGCAGGATCCGACTCGAGTGTCGACTCACAATGCCCCCTTCTTGAGC |
| **oHammy30F** | ATGCGGAGGAATCACTTCCATATGAGCAAGCCCATGCTGCTGA |
| **oHammy30R** | TGCAGGATCCGACTCGAGTGTCGACTCACGCCGCCCGTGCGCGCAT |
| **oHammy31F** | ATGCGGAGGAATCACTTCCATATGAGCAAAACCGTTGAGAACAT |
| **oHammy31R** | TGCAGGATCCGACTCGAGTGTCGACTCACGTCGTCGCCGCCGTGTG |
| **oHammy32F** | ATGCGGAGGAATCACTTCCATATGAGCGGCGAGTCGGCACTA |
| **oHammy32R** | TGCAGGATCCGACTCGAGTGTCGACTCACGCCTGCACCATCAGCGA |
| **oHammy33F** | ATGCGGAGGAATCACTTCCATATGAGCCTGGCTGACCGTCTC |
| **oHammy33R** | TGCAGGATCCGACTCGAGTGTCGACTCAGACAGAGACACGGGCGCC |
| **oHammy34F** | ATGCGGAGGAATCACTTCCATATGTCTCTGTCTGATCGCCTC |
| **oHammy34R** | TGCAGGATCCGACTCGAGTGTCGACTCACAGCACGCCCAATCGGGA |
| **oHammy35F** | ATGCGGAGGAATCACTTCCATATGATGCGGTGCGCTGATCGG |
| **oHammy35R** | TGCAGGATCCGACTCGAGTGTCGACTCAGCTGCCGCGGGTCGCCTG |
| **oHammy36F** | ATGCGGAGGAATCACTTCCATATGAGCACGCCCACAATGACA |
| **oHammy36R** | TGCAGGATCCGACTCGAGTGTCGACTCACTTCGCCCGGTCCTGCTT |
| **oHammy37F** | ATGCGGAGGAATCACTTCCATATGAGCGACGCCGCCGCGGTC |
| **oHammy37R** | TGCAGGATCCGACTCGAGTGTCGACTCATGGCGCAACCACGTCCCG |
| **oHammy38F** | ATGCGGAGGAATCACTTCCATATGACGCTGCAGCGTAAGCCC |
| **oHammy38R** | TGCAGGATCCGACTCGAGTGTCGACTCACTTCGGCGAACGCTTCGCGT |
| **oHammy39F** | ATGCGGAGGAATCACTTCCATATGCAAGAGCACTTTTACCTCGG |
| **oHammy39R** | TGCAGGATCCGACTCGAGTGTCGACTCAGGCGACGTCGAGGTCGAACA |
| **oHammy40F** | ATGCGGAGGAATCACTTCCATATGCGCCTGCGGCCGGGCCGC |
| **oHammy40R** | TGCAGGATCCGACTCGAGTGTCGACTCAGCGGCCGTCGAGCGCCCG |

| | |
|---|---|
| **oHammy41F** | ATGCGGAGGAATCACTTCCATATGCACCCAAAGGTGTACCCA |
| **oHammy41R** | TGCAGGATCCGACTCGAGTGTCGACTCAGGCCCCGAGGTCGTCGGC |
| **oHammy42F** | ATGCGGAGGAATCACTTCCATATGGTGGCAAAAGCTCGGCGC |
| **oHammy42R** | TGCAGGATCCGACTCGAGTGTCGACTCACGCCGCCAATAGGTCGCC |
| **oHammy43F** | ATGCGGAGGAATCACTTCCATATGCGCAAATTGTCGGGCTCG |
| **oHammy43R** | TGCAGGATCCGACTCGAGTGTCGACTCAGTCGGGCTCAGCCTCGGGTG |
| **oHammy44F** | ATGCGGAGGAATCACTTCCATATGACCGTCCTACTGTTTGCC |
| **oHammy44R** | TGCAGGATCCGACTCGAGTGTCGACTCAGCTGACCGGCTCGGGCGGCTG |
| **oHammy45F** | ATGCGGAGGAATCACTTCCATATGGACGACAGCAACAAGTCGCT |
| **oHammy45R** | TGCAGGATCCGACTCGAGTGTCGACTCACAGCCCCGAGACGCCGTTGC |
| **oHammy46F** | ATGCGGAGGAATCACTTCCATATGGGCGACAACGGAATCCGCG |
| **oHammy46R** | TGCAGGATCCGACTCGAGTGTCGACTCACGCGGCGGCCTCGGCAAG |
| **oHammy47F** | ATGCGGAGGAATCACTTCCATATGATCCCGCCGGTTGTCGTC |
| **oHammy47R** | TGCAGGATCCGACTCGAGTGTCGACTCAGGCCCCTAACTGGTCGAC |
| **oHammy48F** | ATGCGGAGGAATCACTTCCATATGTCACTTGCACGCCATATGG |
| **oHammy48R** | TGCAGGATCCGACTCGAGTGTCGACTCAACCCGCGCGATCCATGTCG |
| **oHammy49F** | ATGCGGAGGAATCACTTCCATATGCCGTGCAAAATTCTACGTC |
| **oHammy49R** | TGCAGGATCCGACTCGAGTGTCGACTCACGACACCGCCCGCGCGAG |
| **oHammy50F** | ATGCGGAGGAATCACTTCCATATGAGCCCCGCACCGTACTGC |
| **oHammy50R** | TGCAGGATCCGACTCGAGTGTCGACTCACAGCGACCACACCCCGAC |
| **oHammy51F** | ATGCGGAGGAATCACTTCCATATGAGCTTCTGCATTGCCTACGG |
| **oHammy51R** | TGCAGGATCCGACTCGAGTGTCGACTCACGCCTGCACCGCCTGCCGTG |
| **oHammy52F** | ATGCGGAGGAATCACTTCCATATGAGGGCTGCACAGATCGACGA |
| **oHammy52R** | TGCAGGATCCGACTCGAGTGTCGACTCACTTGTCGGCCTCGCTTTCGG |
| **oHammy53F** | ATGCGGAGGAATCACTTCCATATGAACGCCCTTACCCTGCCAGA |
| **oHammy53R** | TGCAGGATCCGACTCGAGTGTCGACTCATGCGGCGGCCCCCTCACG |
| **oHammy54F** | ATGCGGAGGAATCACTTCCATATGAGCGATCAGCCAATGTGC |
| **oHammy54R** | TGCAGGATCCGACTCGAGTGTCGACTCATCCCTTTGTTTCGTTCGCAG |
| **oHammy55F** | ATGCGGAGGAATCACTTCCATATGCACGCATTCATCAAGGCC |
| **oHammy55R** | TGCAGGATCCGACTCGAGTGTCGACTCACGGGCGATCGACGTCGGG |
| **oHammy56F** | ATGCGGAGGAATCACTTCCATATGACCGACAGCAAGCGCCCG |
| **oHammy56R** | TGCAGGATCCGACTCGAGTGTCGACTCAGGCATTGCTTGCCCTTTCG |
| **oHammy57F** | ATGCGGAGGAATCACTTCCATATGCCTGACATTTCAGAGGTC |
| **oHammy57R** | TGCAGGATCCGACTCGAGTGTCGACTCATCCCTTGTCCGTGTCGACCAG |
| **oHammy58F** | ATGCGGAGGAATCACTTCCATATGGCAAGGCAACTGATTGTCGTC |
| **oHammy58R** | TGCAGGATCCGACTCGAGTGTCGACTCACAGGGCCGCCGCCTTCGT |
| **oHammy59F** | ATGCGGAGGAATCACTTCCATATGAGCAATTTCGCGCACGTG |
| **oHammy59R** | TGCAGGATCCGACTCGAGTGTCGACTCATCGCTGCGCACCGCCCAT |
| **oHammy60F** | ATGCGGAGGAATCACTTCCATATGAGCGGCAACGCAGGCTTT |

| | |
|---|---|
| **oHammy60R** | TGCAGGATCCGACTCGAGTGTCGACTCACTTGGCGCCGCCCTCGCA |
| **oHammy61F** | ATGCGGAGGAATCACTTCCATATGAGCCACTACATGCGCACG |
| **oHammy61R** | TGCAGGATCCGACTCGAGTGTCGACTCAGACGACTACAACGCCGAA |
| **oHammy62F** | ATGCGGAGGAATCACTTCCATATGCAAGATGACAGCCCCCGA |
| **oHammy62R** | TGCAGGATCCGACTCGAGTGTCGACTCACGGCTGCGCCTCGTCGAG |
| **oHammy63F** | ATGCGGAGGAATCACTTCCATATGAGCGCCCGGTCGACGTTC |
| **oHammy63R** | TGCAGGATCCGACTCGAGTGTCGACTCACGGGCATTCACCTTTCGG |
| **oHammy64F** | ATGCGGAGGAATCACTTCCATATGAGCCGTCGGCACAACTGC |
| **oHammy64R** | TGCAGGATCCGACTCGAGTGTCGACTCAGGTCCCGTCGTAGTAGTCGG |
| **oHammy65F** | ATGCGGAGGAATCACTTCCATATGAGCAACGATTCGTACGGATT |
| **oHammy65R** | TGCAGGATCCGACTCGAGTGTCGACTCACTTGACCATGCCGAGCTTCT |
| **oHammy66F** | ATGCGGAGGAATCACTTCCATATGCTGACCGTCTACACGACCGG |
| **oHammy66R** | TGCAGGATCCGACTCGAGTGTCGACTCAGCGGGCCTCGATCGCGGC |
| **oHammy67F** | ATGCGGAGGAATCACTTCCATATGCCCGTCGATACCCGGCTG |
| **oHammy67R** | TGCAGGATCCGACTCGAGTGTCGACTCACAGTGCAAACAGCCCATC |
| **oHammy68F** | ATGCGGAGGAATCACTTCCATATGAACGGCCTAACTGACCTGC |
| **oHammy68R** | TGCAGGATCCGACTCGAGTGTCGACTCACCACCGCGGGGCCTCGTTC |
| **oHammy69F** | ATGCGGAGGAATCACTTCCATATGACTGACCACACTCTCGACC |
| **oHammy69R** | TGCAGGATCCGACTCGAGTGTCGACTCAGGCATTAGCCGCCCGCCAT |
| **oHammy70F** | ATGCGGAGGAATCACTTCCATATGCGCACTCGTTTTGAGGCCC |
| **oHammy70R** | TGCAGGATCCGACTCGAGTGTCGACTCACGACGGCCGCACCTCCGTG |
| **oHammy71F** | ATGCGGAGGAATCACTTCCATATGACGAGTTTGCCGGAAATCG |
| **oHammy71R** | TGCAGGATCCGACTCGAGTGTCGACTCATCCCGTGTCCGAAACTGACGG |
| **oHammy72F** | ATGCGGAGGAATCACTTCCATATGCCTTTGAAGCGCAACAGATTA |
| **oHammy72R** | TGCAGGATCCGACTCGAGTGTCGACTCACTTCGCGCCCTCGTTCAGGTG |
| **oHammy73F** | ATGCGGAGGAATCACTTCCATATGAAGCGCGTCAAGACTGTTC |
| **oHammy73R** | TGCAGGATCCGACTCGAGTGTCGACTCACGCTGCGGCGTTCGCGCGG |
| **oHammy74F** | ATGCGGAGGAATCACTTCCATATGAACCGCAATCTCATTCTCG |
| **oHammy74R** | TGCAGGATCCGACTCGAGTGTCGACTCAGTCGCTGGCACCTCCCCCG |
| **oHammy75F** | ATGCGGAGGAATCACTTCCATATGAGCGAAGTCATCGACTACAG |
| **oHammy75R** | TGCAGGATCCGACTCGAGTGTCGACTCAGCGAGTTGCGTACTTGCG |
| **oHammy76F** | ATGCGGAGGAATCACTTCCATATGCACAACACCCACGTTTAC |
| **oHammy76R** | TGCAGGATCCGACTCGAGTGTCGACTCAGTCGGCCGCCATGTGCGCCA |
| **oHammy77F** | ATGCGGAGGAATCACTTCCATATGACCGAAAACATCACCCGC |
| **oHammy77R** | TGCAGGATCCGACTCGAGTGTCGACTCAGGACTCGATGCGGTCGTG |
| **oHammy78F** | ATGCGGAGGAATCACTTCCATATGAACACGAAAGATCCGAGG |
| **oHammy78R** | TGCAGGATCCGACTCGAGTGTCGACTCACACCGCCACCGCCTCGAC |

| | |
|---|---|
| **oHammy79F** | ATGCGGAGGAATCACTTCCATATGAGCGCACTCTCTGTCGCT |
| **oHammy79R** | TGCAGGATCCGACTCGAGTGTCGACTCACCAGAGCGACCCCAGCCC |
| **oHammy80F** | ATGCGGAGGAATCACTTCCATATGATCGCGCTCACTGAAATGC |
| **oHammy80R** | TGCAGGATCCGACTCGAGTGTCGACTCACATCACGCACACATAGCCG |
| **oHammy81F** | ATGCGGAGGAATCACTTCCATATGATGCTGTCCGTTGAGCCGG |
| **oHammy81R** | TGCAGGATCCGACTCGAGTGTCGACTCATTTCGGGTCGGGTCGGTCA |
| **oHammy82F** | ATGCGGAGGAATCACTTCCATATGCGAGGACTGATCGACAGG |
| **oHammy82R** | TGCAGGATCCGACTCGAGTGTCGACTCAGCCGATACAGTCGCCGCA |
| **oHammy83F** | ATGCGGAGGAATCACTTCCATATGTTGGCACCTCAACAGATCAG |
| **oHammy83R** | TGCAGGATCCGACTCGAGTGTCGACTCACTGCCCCTTCACGTTCAAG |
| **oHammy84F** | ATGCGGAGGAATCACTTCCATATGAGCACAGAAGGCTTTTCGC |
| **oHammy84R** | TGCAGGATCCGACTCGAGTGTCGACTCACCCATGCCGCAACGCCTGC |
| **oHammy85F** | ATGCGGAGGAATCACTTCCATATGGGTGAGTCGGCGCTGTTC |
| **oHammy85R** | TGCAGGATCCGACTCGAGTGTCGACTCACGCTGCCACCGCCGCAGG |
| **oHammy86F** | ATGCGGAGGAATCACTTCCATATGAGCGGCGATCTGCGCGAG |
| **oHammy86R** | TGCAGGATCCGACTCGAGTGTCGACTCATCGCCTCGCCCCCGTCCC |
| **oHammy87F** | ATGCGGAGGAATCACTTCCAATGAGCGTCGCGGTTGTGGCG |
| **oHammy87R** | TGCAGGATCCGACTCGAGTGTCGACTCAGAACGGTGGCGGCAAGCT |
| **oHammy88F** | ATGCGGAGGAATCACTTCCATATGCAGGAATACACCAAGGCC |
| **oHammy88R** | TGCAGGATCCGACTCGAGTGTCGACTCACTTGCCCTTGGGCGCCTTG |
| **oHammy89F** | ATGCGGAGGAATCACTTCCATATGCAGAACACAATCGCCGCC |
| **oHammy89R** | TGCAGGATCCGACTCGAGTGTCGACTCAGGCGATTTCTGACCACTC |
| **oHammy90F** | ATGCGGAGGAATCACTTCCATATGACCGACACGATTCACGCA |
| **oHammy90R** | TGCAGGATCCGACTCGAGTGTCGACTCAGGCCGCCGCGGCGCGCCG |
| **oHammy91F** | ATGCGGAGGAATCACTTCCATATGTACGCATGGATCAATGGTCAG |
| **oHammy91R** | TGCAGGATCCGACTCGAGTGTCGACTCAGCAGCCGTGCAGCATCTTG |
| **oHammy92F** | ATGCGGAGGAATCACTTCCATATGAGCACGACCGTTACCTACC |
| **oHammy92R** | TGCAGGATCCGACTCGAGTGTCGACTCAGGCCGCCGCGGGCTCGAC |
| **oHammy93F** | ATGCGGAGGAATCACTTCCATATGGCTCTAATCAGCCCTGAG |
| **oHammy93R** | TGCAGGATCCGACTCGAGTGTCGACTCAGAGCTTGAAGGGGAAGGGC |
| **oHammy94F** | ATGCGGAGGAATCACTTCCATATGGCACAAGACAGCACCGAG |
| **oHammy94R** | TGCAGGATCCGACTCGAGTGTCGACTCAGTCGAGCGGTTCGTACTT |
| **oHammy95F** | ATGCGGAGGAATCACTTCCATATGAACGAAGCTCGCAACACTG |
| **oHammy95R** | TGCAGGATCCGACTCGAGTGTCGACTCAGGGCCGCCAGACCCGAGCG |
| **oHammy9_internal1** | GGTCACGATCACTGCAAGTGC |
| **oHammy9_internal2** | GAAACCGCTACGCGCG |
| **oHammy20_internal1** | CTGCCTCTGCGGTGGC |

| | |
|---|---|
| **oHammy20_internal2** | CGACTGGTCGACAAGGTGC |
| **oHammy20_internal3** | CACCTTGTCGTGAGCCCG |
| **oHammy20_internal4** | GTCGTTCGGCCGGTAACC |
| **oHammy22_internal1** | CGATCCTGGCCTTGCCAAC |
| **oHammy26_internal1** | CAGCCAGCGGCTCGATC |
| **oHammy26_internal2** | GACACGTTGCCGGTATCCG |
| **oHammy26_internal3** | CGATCGGGCAGCAGCAAG |
| **oHammy29_internal1** | CGTTCGACCGTTACGCC |
| **oHammy68_internal1** | CGTCACGATCGACGG |
| **oHammy68_internal2** | GGTTGCCCATCAAGAACAGC |
| **oHammy68_internal3** | GGTCACGTCGCAACCAAG |
| **pExTra_seqF** | GTACCCGTGTGTACGACCAGC |
| **pExTra_universalR** | CCCTTCGAGACCATAGATCTGTTCC |
